# Administration of barcoded AAV capsid library to the putamen of non-human primates identifies variants with efficient retrograde transport

**DOI:** 10.1101/2025.07.07.663544

**Authors:** Yulia Dzhashiashvili, Jodi L. McBride, Emily Fabyanic, Xin Huang, Brian M. Kelly, Greglynn D. Walton-Gibbs, Mohamad Nayal, Ariel A. Hippen, Zhenming Yu, Pichai Raman, Elizabeth Ramsburg, Marcus Davidsson, Esteban A. Engel, Tomas Björklund

## Abstract

Adeno-associated viral vectors have become a leading choice for gene therapy in the central nervous system due to their safety profile, efficient neuronal transduction, and capacity for sustained transgene expression. We previously reported that AAV2-derived capsids developed using the BRAVE (Barcoded Rational AAV Vector Evolution) approach have enhanced retrograde transport properties in the rodent brain, compared to parental AAV2. Retrograde transport enables broader coverage of connected brain regions after a single focal intraparenchymal brain injection and is therefore a powerful tool for delivery of vectors to distant sites with potentially higher specificity, transduction efficacy and safety. Because transport properties can vary among species, we further characterized a barcoded library of 25 BRAVE-derived AAV2 capsid variants, along with the parental AAV2 serotype and benchmark AAV capsids, in brains of adult cynomolgus monkeys after intraputaminal dosing. Based on RNA and DNA amplicon sequencing, single-nucleus RNA sequencing, and histological assessment, we report here capsid variants with enhanced retrograde transport and expression compared to the parental AAV2 capsid. These properties make them potentially useful for disease indications in which broader brain coverage is desirable beyond the injection site.

## INTRODUCTION

A recombinant adeno-associated virus (rAAV) is a modified, replication-defective, and non-pathogenic version of the naturally occurring AAV, where the viral DNA is replaced with a therapeutic transgene.^1–3^ rAAV-based therapies have shown promise in preclinical and clinical studies to treat various central nervous system (CNS) disorders.^4,5^ Direct brain administration of rAAVs via stereotaxic intraparenchymal (IPa) injection is a widely used gene therapy delivery strategy.^6^ It allows for bypassing the blood-brain barrier and requires only small quantities of vector to precisely deliver the gene of interest. Moreover, the retrograde axonal transport property of AAVs, which is serotype-dependent, can be harnessed for therapeutic purposes to transduce functionally connected brain regions distal to the IPa injection site. In many CNS disorders, disease-specific pathologies extend beyond the initially affected brain region, impacting distributed circuits and networks.^7–10^ Thus, it is desirable to engineer AAV capsids with both high retrograde transport efficiency and specific cell tropism, as vehicles to deliver AAVs throughout the brain and to disease-affected cell types.

Davidsson et al.^11^ have previously described the BRAVE (Barcoded Rational AAV Vector Evolution) approach to design and screen an AAV2-based library with the aim to develop AAV capsid variants that are efficiently taken up at axonal terminals and transported retrogradely in neurons *in vivo*. The BRAVE-derived library of MNM001-MNM025 AAV capsid variants was characterized in the rodent brain, in human embryonic stem cell-derived dopaminergic (DA) neurons *in vitro* and in the brain of rats with human-derived DA transplants.^11^ It was shown that the herpes simplex virus type 2 (HSV-2)-derived peptide sequence used in MNM004 enabled efficient retrograde transport to all afferent rat brain regions following striatal injection, which is a robust improvement over the parental AAV2 capsid. It was also shown that the canine adenovirus type 2 (CAV-2)-derived peptide sequence used in MNM008 significantly improved retrograde spread to nigral neurons compared to parental AAV2.^11^ The properties of these AAV2 variants make them potentially amenable for use in future clinical gene therapies.

To further investigate the properties and potential translational efficacy of BRAVE-derived AAV2 variants, we conducted a study in nonhuman primates (NHPs), taking advantage of their highly developed brain structure and neuroanatomical pathways, which are similar to the human brain.^12^ In this study, we evaluated a library of 25 barcoded AAV2 variants (i.e., MNM001-MNM025) in 3 adult cynomolgus monkeys following bilateral IPa administration into the putamen. These 25 AAV variants were compared with the parental AAV2 capsid and benchmark capsids AAV2-retro,^13^ AAV9, and AAV9-retro.^14^ Multiple samples from 15 target brain regions were analyzed by sequencing of DNA barcodes to assess vector genome biodistribution and by sequencing of RNA-expressed barcodes to identify top functional capsids. Furthermore, additional brain samples were processed via a modified snRNAseq method to investigate the cell-type tropism of barcoded capsid variants. Our screening results established MNM004 as a cross-species compatible capsid with dramatically improved retrograde transport properties over parental AAV2. We also identified MNM021 as a capsid with enhanced retrograde trafficking to multiple brain structures, while retaining robust expression at the injection site in the putamen. Overall, the findings of this study will help guide the selection of the best retrograde AAV capsid for treatment of specific brain diseases.

## RESULTS

### Characterization of the AAV capsid library

Each MNM001-MNM025 AAV capsid variant displays a unique peptide, derived from a known neuron-related protein from the following categories: neurotropic viruses, lectins, neurotrophins, neurotoxins, and neuronal proteins (**Table S2**).^11^ These unique polypeptide sequences were inserted into the AAV2-Cap gene at position N587,^15^ thereby mutating the wild-type (WT) AAV2 heparan sulfate proteoglycan (HSPG)-binding motif. HSPGs are the primary cellular receptors for WT AAV2.^16^ Additionally, the AAV genome plasmid contains a unique 20 base pair (bp) molecular barcode, inserted downstream of the enhanced green fluorescent protein (EGFP) reporter gene in the 3’ untranslated region (UTR), linking the genome to its respective peptide (**Figure 1A**). We analyzed a library of 29 AAV capsids (**Table S1**). The capsid variants were packaged individually to eliminate the risk of chimeras or recombination. Prior to injection into NHPs, an aliquot of the complete AAV vector library was sequenced. The results showed balanced capsid abundance in the input library, with the average capsid variant abundance of 3.2%. All downstream barcode amplicon sequencing data (DNA, RNA, snRNA amplicon) were normalized to the relative AAV library input abundance for each capsid.

**Figure 1.**
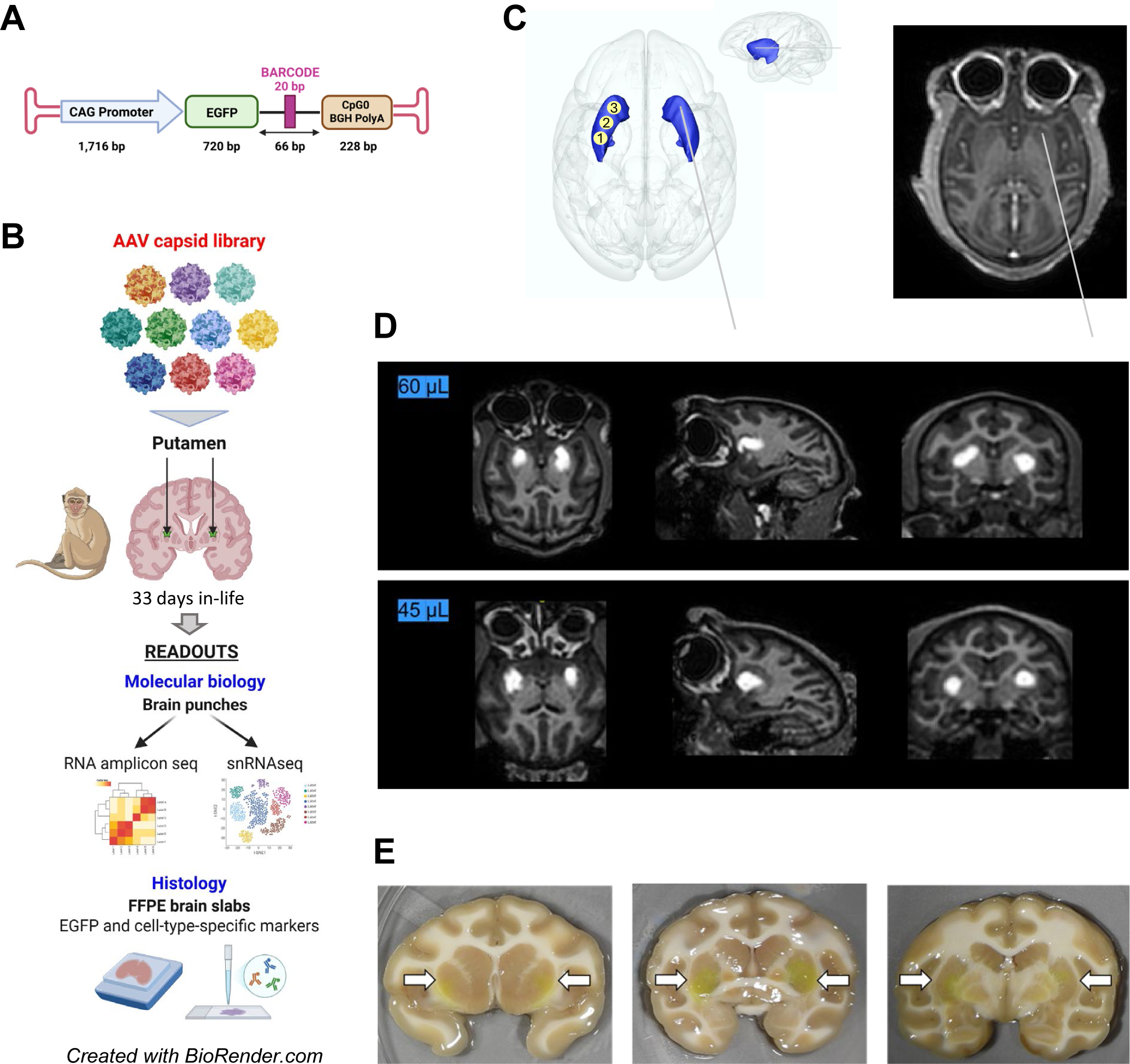
Barcoded AAV capsid library, experimental strategy, and surgical delivery. (A) Representative barcoded AAV genome construct. Each capsid variant contains a CAG-EGFP expression cassette with a unique 20 bp barcode positioned between the EGFP reporter gene and CpG0 BGH PolyA tail. The rAAV plasmid length is 10 kb; ITR-to-ITR length is 3 kb. (B) Schematic overview of the experimental strategy. Note that molecular biology readouts also include DNA amplicon seq (omitted from the schematic for clarity). (C) The diagram of surgical methodology shows AAV delivery at 3 locations along the axis of the putamen with Clearpoint Smartflow^®^ Neuro Cannula, using CED and occipital trajectories. (D) Comparison of infusate spread by injection volume. T1-weighted representative images obtained approximately 15 min after Gadoteridol-containing test article administration at 60 µL and 45 µL injection volumes per hemisphere. Scans were performed using a 3T Philips Achieva MRI scanner. (E) Visible EGFP protein expression in fresh coronal brain slabs by unaided eye. White arrows point to the putamen (injection site, bilaterally). Notably, there was no edema, discoloration, or distortion of the brain tissue. rAAV, recombinant AAV; BGH PolyA, bovine growth hormone polyadenylation signal; EGFP, enhanced green fluorescent protein; CED, convection enhanced delivery; MRI, magnetic resonance imaging.

### Study design

Cynomolgus monkeys (*Macaca fascicularis*) were selected as a large preclinical animal model for the simultaneous analysis of AAV vector biodistribution and transgene expression in the brain. NHPs are the closest to humans in terms of anatomical brain structure and neural connectivity, as well as physiology and immunity.^12^ Animals were selected for the study based on their AAV-neutralizing antibody (NAb) status to AAV2 and AAV9 capsids. All animals enrolled in the study exhibited serum NAb titers of either <1:1 or 1:1, indicating no pre-existing neutralizing antibodies or minimal levels of pre-existing neutralizing antibodies to AAV2 and AAV9. Animals were also confirmed to be seronegative to HSV-1 and HSV-2. The choice of the putamen as a brain region to evaluate the AAV library biodistribution was driven by the following considerations: 1) the putamen is affected by a number of pathological and neurodegenerative conditions potentially amenable to gene therapy^17,18^; 2) the putamen afferents are very well established and can be utilized for retrograde AAV delivery to distal brain structures^19^; 3) the putamen is a large structure in the forebrain and is therefore easily accessible with potentially fewer surgical complications. Animals were monitored for approximately 1-month post-dosing (Day 33 ± 1). The study design is shown in **Table 1** and the schematic experimental workflow is detailed in **Figure 1B**.

**Table 1.** NHP study design: Group designation and dose level.

| Group | Animal Number | Test Article | Dose Level (vg/brain)<sup>a</sup> | Concentration (vg/mL) | Volume per Hemisphere (mL) | Necropsy Time Point |
| --- | --- | --- | --- | --- | --- | --- |
| 1 | 1501 | AAV Library | 1.0E12 | 8.33E12<sup>b</sup> | 0.060 | Study Day 33 ± 1 |
|  | 1001 |  | 0.75E12 |  | 0.045 |  |
|  | 1002 |  |  |  |  |  |

All animals received 20 mg methylprednisolone via intramuscular administration once on Study Day 1, prior to AAV administration.
<sup>a</sup> Following test article administration in animal 1501, a decision was made to lower the infusion volume from 60 µL to 45 µL per hemisphere in the remaining 2 animals.
<sup>b</sup> Test article contains 2 mM ProHance (Gadoteridol), a gadolinium-based contrast reagent to visualize vector spread.
AAV, adeno-associated virus; vg, vector genome.

### Volumetric analysis of gadoteridol coverage

Vector was delivered into the bilateral putamen using intra-MRI guidance and the SmartFlow® Neuro Cannula^20^ (ClearPoint Neuro; Solano Beach, CA). An occipital trajectory to the putamen was selected to maximize distribution throughout the structure, and AAV vector was injected in three separate deposits along the axis of the putamen using convection enhanced delivery (CED)^21^ and a slow-ramping infusion rate (**Figure 1C**). In the first surgical case, the animal was injected into the putamen with 60 µL of vector per hemisphere (20 µL per deposit) (**Figure 1D, top panel**). Gadoteridol was primarily localized to the putamen; however, minor diffusion was observed outside of the boundaries of the structure. As the goal was to maintain the spread of the vector within the borders of the putamen, the infusion volume was lowered to 45 µL per hemisphere for the remaining 2 animals (15 µL per deposit), resulting in a more restricted distribution of the vector within the target area (**Figure 1D, bottom panel**). Volumetric analysis showed a 34-49% coverage of the left putamen (42 ± 8%) and a 31-54% coverage of the right putamen (42 ± 12%) (**Table 2**). At necropsy, 4 mm thick brain slabs collected through the putamen showed evidence of EGFP that was visible upon macroscopic investigation (**Figure 1E**), confirming appropriate intraputaminal targeting.

**Table 2.** Volumetric analysis of Gadoteridol coverage in the putamen.

| Animal ID | Injection Volume (µL) | Left Putamen Volume (mm<sup>3</sup>) | Left Gad Volume (mm<sup>3</sup>) | Percent Coverage Left Hem | Right Putamen Volume (mm<sup>3</sup>) | Right Gad Volume (mm<sup>3</sup>) | Percent Coverage Right Hem |
| --- | --- | --- | --- | --- | --- | --- | --- |
| 1501 | 60 | 510.3 | 250.0 | 49% | 526.0 | 283.0 | 54% |
| 1002 | 45 | 530.0 | 181.4 | 34% | 545.8 | 218.8 | 40% |
| 1001 | 45 | 505.0 | 217.0 | 43% | 510.0 | 160.0 | 31% |

Gad, gadoteridol; Hem, hemisphere; ID, identification.

### The MNM004 capsid promotes highly efficient retrograde transport

A total of 29 capsids were analyzed in 3 NHPs. Results are shown for 3 select capsids: parental AAV2, AAV2-retro, and MNM004 (**Figure 2, Figure S1**). Multiple brain regions were profiled, including 7 cortical areas (dorsal prefrontal cortex, ventral prefrontal cortex, anterior cingulate cortex, dorsal premotor cortex, ventral premotor cortex, primary motor cortex, and somatosensory cortex), the basal ganglia structures (caudate, putamen, external and internal globus pallidus), thalamus, hippocampus, and amygdala. Both DNA and RNA barcode amplicon sequencing data showed strong evidence of animal-to-animal consistency (**Figure 2**). The data demonstrated that AAV2-retro (peptide insertion LADQDYTKTA) and MNM004 (peptide insertion VMSVLLVDTDATQQ) emerged as leading retrograde capsids, with markedly improved biodistribution and expression over parental AAV2 (**Figure 2, Figure S1**). These two capsids were abundantly expressed throughout all cortical areas profiled, as well as in the amygdala. Interestingly, DNA amplicon sequencing revealed DNA barcode enrichment in cortical areas for AAV2-retro, but not for MNM004 (**Figure 2**).

**Figure 2.**
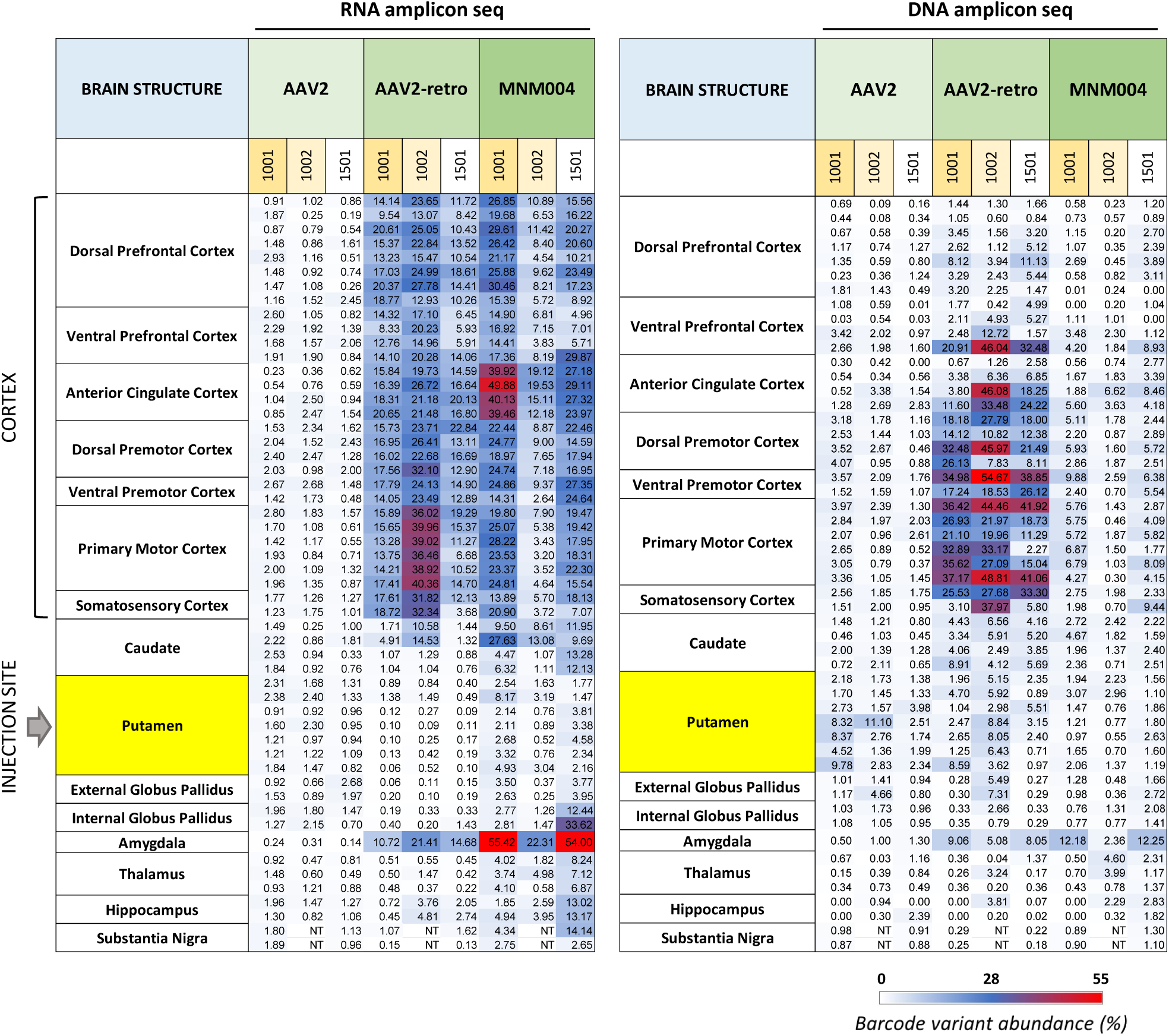
The MNM004 capsid promotes highly efficient retrograde transport. Amplicon sequencing data comparing biodistribution (DNA) and expression (RNA) of select capsids in brain cortical regions and subcortical structures. Data for each capsid (MNM004, parental AAV2, and benchmark AAV2-retro) are shown side-by-side for all 3 NHPs in the study (1001, 1002, and 1501). Except for amygdala, 2 or more samples (punches or dissected areas) per brain structure were collected and analyzed. Each row in the RNA and DNA data tables corresponds to individual brain sample data that show percent capsid barcode variant abundance, out of 100% total for the entire library. NHP, non-human primate; NT, not tested.

### The MNM021 capsid promotes both extensive retrograde transport and robust transduction at the injection site

From the RNA barcode amplicon sequencing screen, capsid MNM021 (peptide insertion DPGYAETPYASVSH) emerged as more widely distributed and abundantly expressed throughout the brain, compared to parental AAV2. Based on relative RNA barcode variant abundance (%), MNM021 was more highly expressed in the putamen, caudate, internal and external globus pallidus, thalamus, and hippocampus. MNM021 also demonstrated enhanced retrograde transport to all cortical regions profiled (**Figure 3, Figure S2, and Figure S3**). It notably differed from MNM004 and AAV2-retro in its comparatively less potent retrograde functionality, i.e. transport to cortical areas, and its lesser presence in the amygdala, where MNM004 and AAV2-retro were both highly expressed (**Figure S2**). Finally, MNM021 was found to be markedly expressed at the injection site, compared to parental AAV2 and to the rest of the capsids in the library (**Figure S2 and Figure S3**).

**Figure 3.**
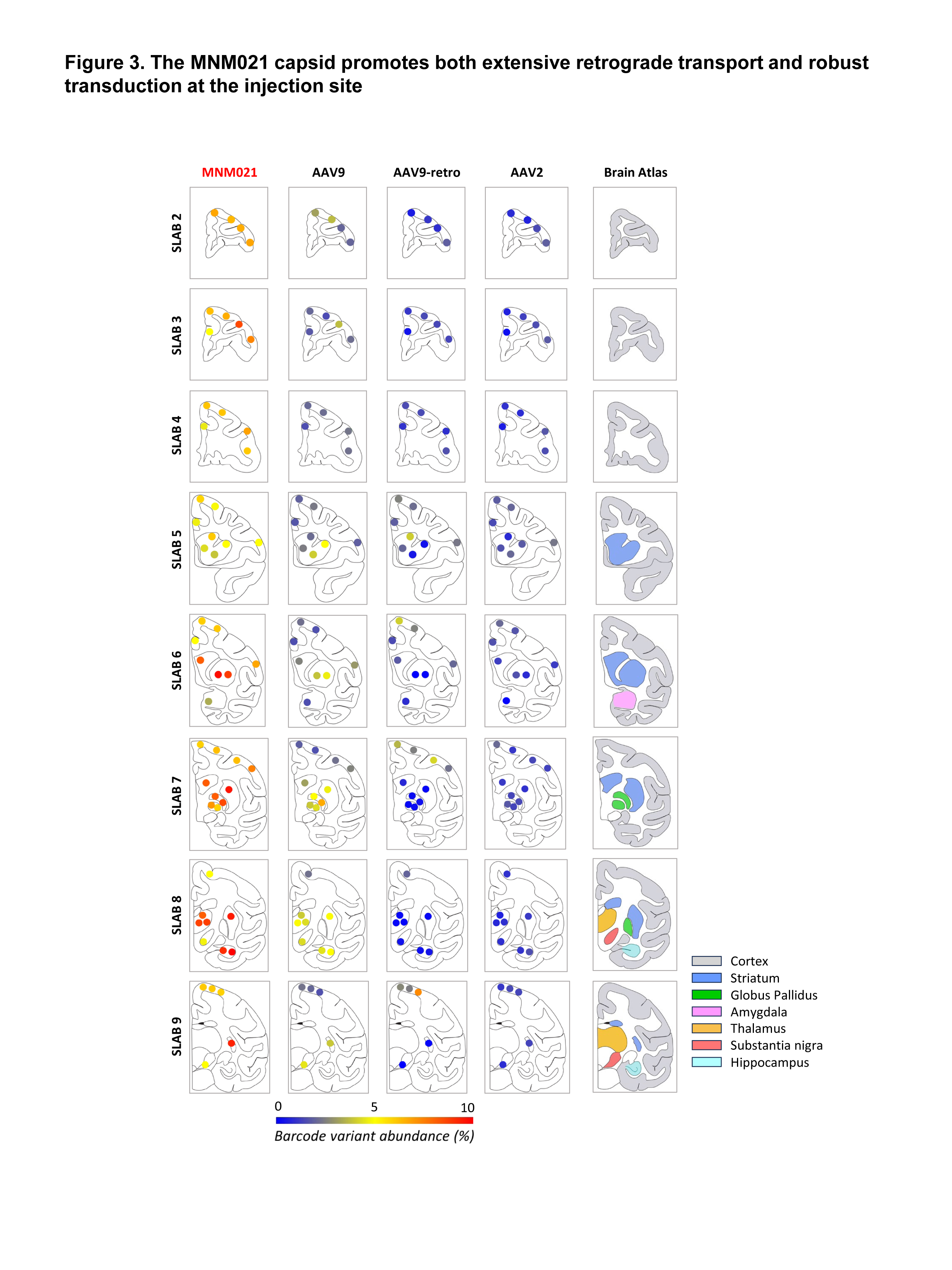
The MNM021 capsid promotes both extensive retrograde transport and robust transduction at the injection site. Visual representation (heatmaps) of RNA amplicon sequencing data for MNM021, parental AAV2 and benchmark AAV9 and AAV9-retro capsids. For each capsid, data were aggregated from 2 NHPs (for substantia nigra) or from 3 NHPs (for all other brain samples). Diagrams demonstrate samples profiled within consecutive rostral-to-caudal coronal brain slabs. For each brain slab, a corresponding anatomical atlas brain map is displayed for reference. The actual numeric NGS data for each brain sample (% barcode variant abundance) were directly and automatically converted into colored pixels with red indicating higher expression values and blue indicating signal values below the threshold. The intensity of the red/blue palette correlates with the numeric values. Brain samples were collected as 3 mm punches, except globus pallidus and substantia nigra, which were microdissected and either split in half (external and internal globus pallidus) or collected as a whole (substantia nigra). For these three structures, colored circles represent data from dissected tissues. NHP, non-human primate; NGS, next-generation sequencing.

### Assessment of other library capsids by RNA and DNA barcode amplicon sequencing

We also profiled the commonly used capsids AAV9 and AAV9-retro (peptide insertion LADQDYTKTA). Interestingly, while AAV9-retro displayed the biodistribution pattern consistent with retrograde trafficking, as expected, it was less prominent compared to AAV2-retro and to the BRAVE variant MNM004 (**Figure S2**). RNA-derived barcode expression results for the rest of the capsids in the library are shown in **Figure S3**. Note that AAV2-retro and MNM004 results were omitted for clarity, as they were significantly more abundant in the cortex compared to the rest of the library and would obscure expression data for other capsids.

### AAV library cell type tropism in injected and distal brain regions

To investigate the cell type-specific tropism of the AAV library capsids in the injected brain region, as well as in brain regions that send afferent projections to the putamen, we evaluated brain punches from the putamen, substantia nigra and anterior cingulate cortex. Tissue from each region was processed for nuclear isolation and snRNAseq, paired with barcode enrichment sequencing. Following quality control and normalization, nuclei were clustered into groups based on their gene expression profiles and visualized using Uniform Manifold Approximation and Projection (UMAP). Cell clusters were manually annotated using prior knowledge of cell type marker genes for each brain region.

### Putamen

For putaminal tissue analysis, cell type clusters included inhibitory medium spiny neurons bearing either DRD1 dopaminergic receptors (MSN1) or DRD2 dopaminergic receptors (MSN2), inhibitory neurons including other MSN subtypes and interneurons, vascular endothelial cells, mature oligodendrocytes, oligodendrocyte progenitor cells (OPCs), astrocytes, microglia, perivascular macrophages (PVMs), T-cells, and a small population of other (uncharacterized) cell types (**Figure 4C and Figure S4**). Based on EGFP mRNA expression, the AAV capsid library overall displayed strong neurotropic properties in the putamen (**Figure 4B**). The MSN1, MSN2, and inhibitory cell clusters had, on average, more than 4 EGFP RNA counts per cell. In contrast, all capsids showed much lower expression (less than 1 EGFP RNA count per cell) in the other cell types. Because EGFP mRNA expression did not reveal the cell type tropism of individual AAV capsids, we relied on the barcode enrichment-based snRNAseq screen to determine percent capsid abundance per cell type (out of 100% total) for neurons and glia (oligodendrocytes, microglia, and astrocytes). The top 8 most abundant capsids are shown in **Figure 4A**, with benchmark AAV9 capsid used as a reference. Notably, the top capsids transducing neurons in the putamen were all AAV2-derived MNM variants, including MNM021. Capsid tropism data were consistent with our histological assessment of EGFP protein expression in coronal brain sections. Immunohistochemical staining for EGFP and the pan neuronal marker NeuN revealed predominantly neuronal tropism within the putamen for all capsids, with most EGFP-positive cells double-positive for NeuN (**Figure 6A**).

**Figure 4.**
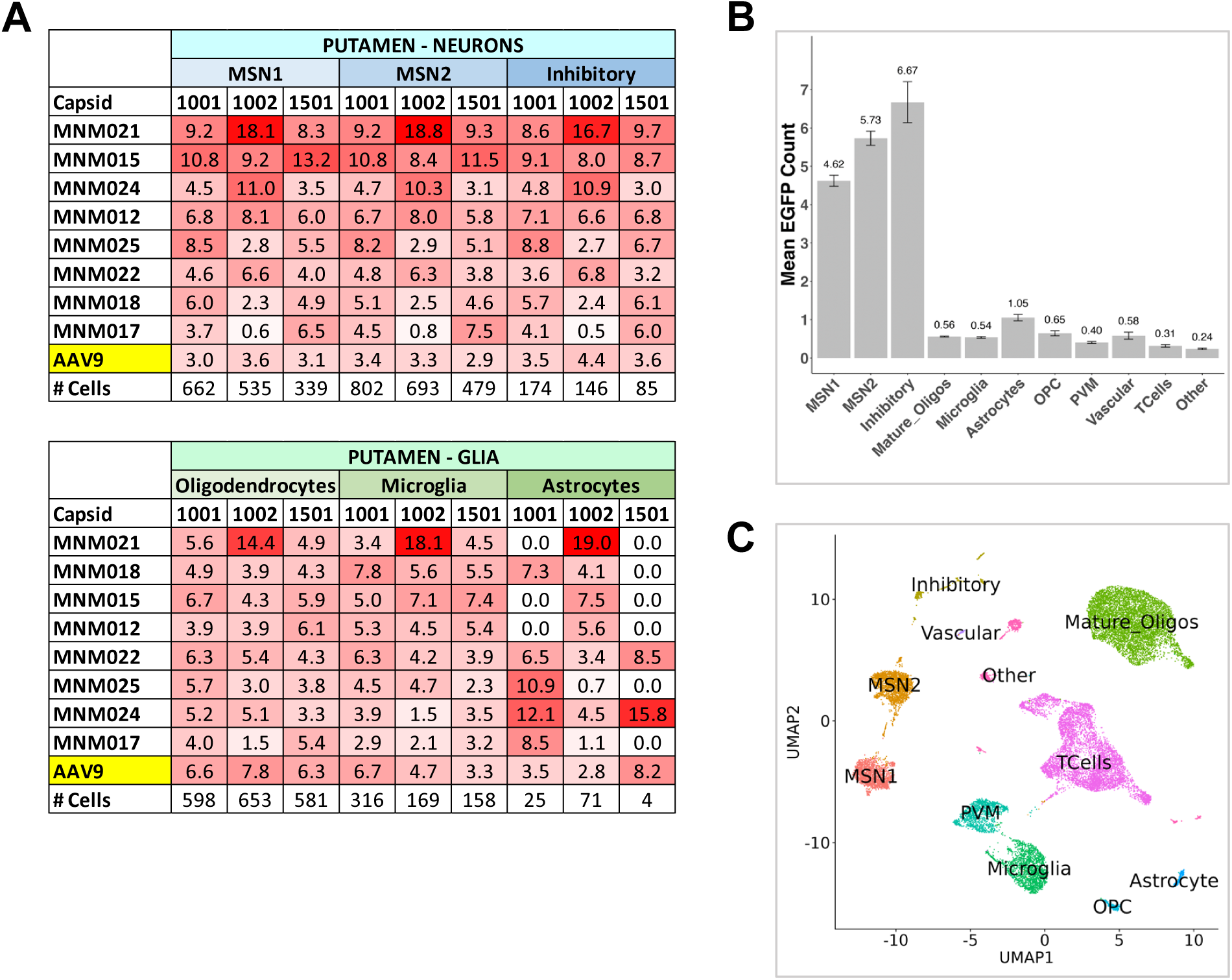
AAV library capsid variant cell-type tropism and EGFP expression in putamen. (A) Barcode enrichment-based snRNAseq data comparing percent capsid variant abundance in neuronal subtypes (MSN1, MSN2, Inhibitory) and glial subtypes (oligodendrocytes, microglia, astrocytes). Data are shown for the top 8 most abundant capsids; benchmark AAV9 capsid was used as a reference. Data for each capsid are shown for all 3 NHPs in the study (1001, 1002, and 1501). #Cells, cell number profiled for each cell subtype per animal. (B) Mean EGFP mRNA count per cell (cluster analysis); data averaged for 3 NHPs. (C) UMAP of cell types identified in the putamen by snRNAseq. MSN1, medium spiny neurons type 1; MSN2, medium spiny neurons type 2; NHP, non-human primate; EGFP, enhanced green fluorescent protein; UMAP, Uniform Manifold Approximation and Projection.

### Substantia Nigra

The AAV library capsids displayed neurotropic properties in the substantia nigra (**Figure 5B**), with preferential transduction of dopaminergic and inhibitory neuronal subtypes. An absence of EGFP mRNA expression in non-neuronal cell types (**Figure 5B**) indicated that the vector reached the substantia nigra pars compacta (SNpc) via axonal transport and not via diffusion from the injection site. The neuronal EGFP mRNA expression findings were concordant with our histological results obtained from coronal brain sections, where we detected robust EGFP protein expression in neurons within the SNpc. To corroborate the cell type-specific tropism, we double-stained brain sections for EGFP and tyrosine hydroxylase (TH), a marker of dopaminergic neurons. We demonstrated that EGFP-positive neurons within the substantia nigra were also positive for TH (**Figures 5D and 6B**). Profiling neurons for capsid variant tropism using snRNAseq proved to be a challenge, as the substantia nigra is known to contain a high percentage of glia: 95% overall, with 72% oligodendrocytes,^22^ which was corroborated by our data (**Figure 5C**). Despite the low number of neuronal nuclei recovered, we identified robust dopaminergic and inhibitory neuronal clusters based on characteristic gene signatures for these neuronal subtypes (**Figure S5**). The barcode enrichment-based snRNA screen revealed MNM021 as the leading capsid variant transducing dopaminergic neurons and inhibitory neurons (**Figure 5A**).

**Figure 5.**
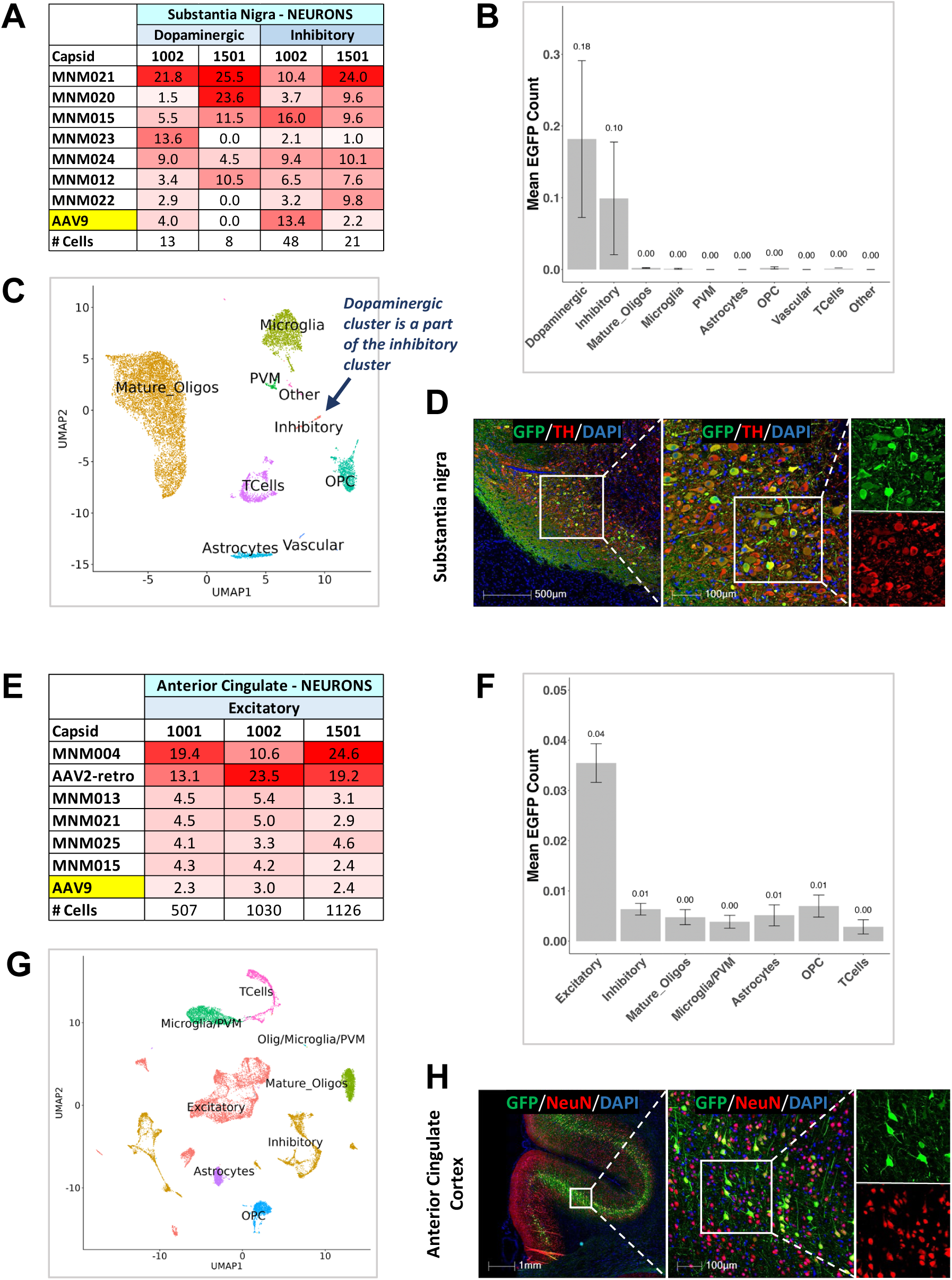
AAV library capsid variant cell-type tropism and EGFP expression in substantia nigra and anterior cingulate cortex. (A-D) Substantia nigra. (A) Barcode enrichment-based snRNAseq data comparing percent capsid variant abundance in neuronal subtypes (dopaminergic, inhibitory) for 2 NHPs (1002 and 1501). Data are shown for the top 7 most abundant capsids; benchmark AAV9 capsid was used as a reference. #Cells, cell number profiled for each cell subtype per animal. (B) Mean EGFP mRNA count per cell (cluster analysis); data averaged for 2 NHPs. (C) UMAP of cell types identified in the substantia nigra by snRNAseq. (D) EGFP protein (green) is expressed in TH-positive (dopaminergic) neurons (red). (E-H) Anterior cingulate. (E) Barcode enrichment-based snRNAseq data comparing percent capsid variant abundance in excitatory neurons for all 3 NHPs in the study (1001, 1002, and 1501). Data are shown for the top 6 most abundant capsids; benchmark AAV9 capsid was used as a reference. #Cells, cell number profiled per animal. (F) Mean EGFP mRNA count per cell (cluster analysis); data averaged for 3 NHPs. (G) UMAP of cell types identified in the anterior cingulate by snRNAseq. (H) EGFP protein (green) is expressed in NeuN-positive neurons (red) within deeper layers of the cortex. EGFP, enhanced green fluorescent protein; NHP, non-human primate; UMAP, Uniform Manifold Approximation and Projection; TH, tyrosine hydroxylase; NeuN, neuronal nuclear antigen.

**Figure 6.**
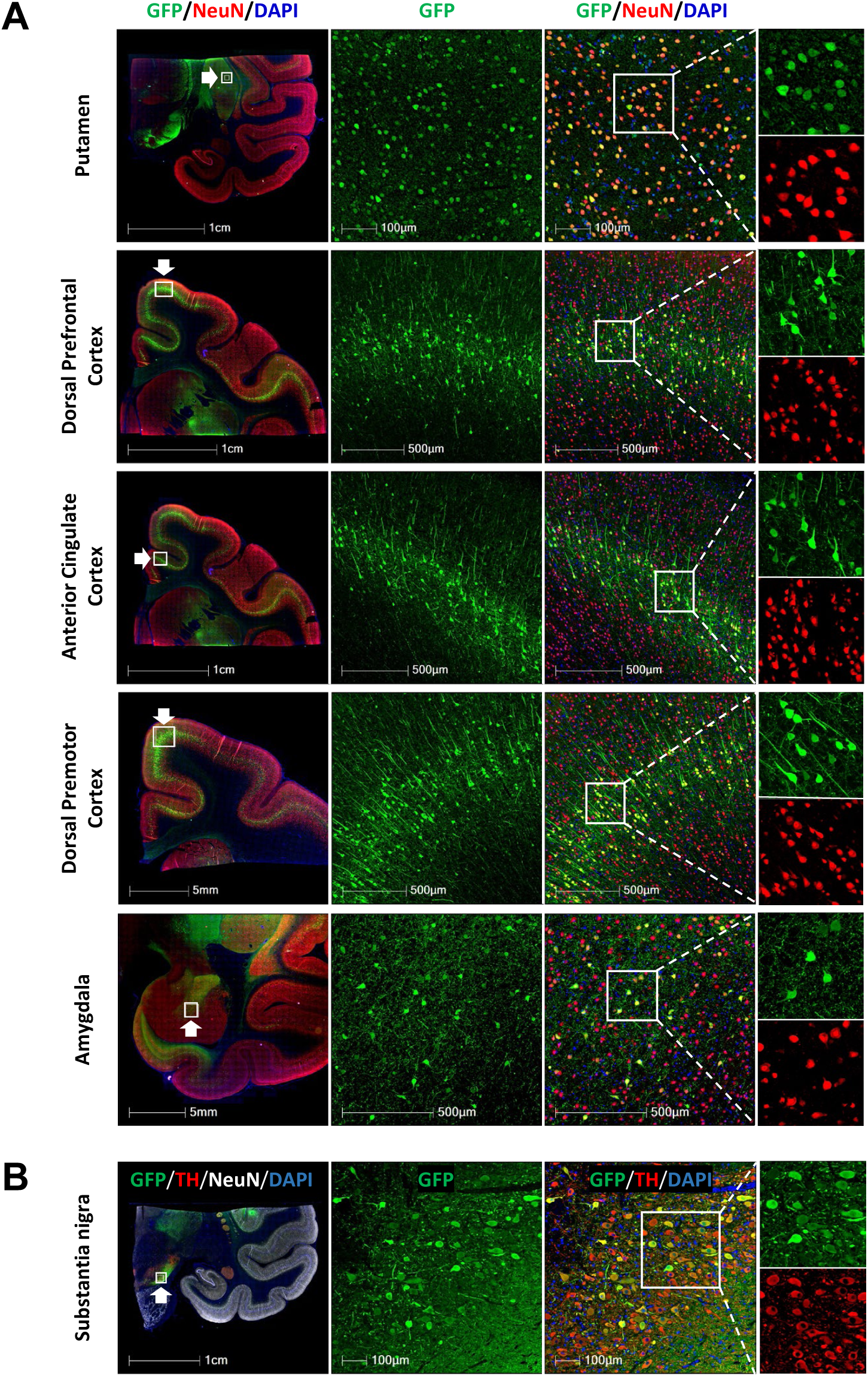
AAV capsid biodistribution by immunohistochemistry. Biodistribution of the AAV capsid library following intraputaminal injection in adult cynomolgus monkeys: axonal transport to distal regions in coronal brain sections. (A) Robust EGFP protein (green) expression at the injection site (putamen), as well as in multiple cortical structures (within deeper cortical layers) and in amygdala, consistent with extensive AAV retrograde transport. Neuronal EGFP protein expression is confirmed by co-staining for NeuN (red), the pan neuronal marker. White arrows point to the regions shown as high-magnification images. (B) Robust EGFP protein (green) expression in substantia nigra pars compacta. EGFP-positive neuronal cell bodies co-express TH (red), the dopaminergic neuronal marker. White arrow points to the region shown as high-magnification images. EGFP, enhanced green fluorescent protein; NeuN, neuronal nuclear antigen; TH, tyrosine hydroxylase.

### Anterior Cingulate Cortex

As in the substantia nigra, the AAV library capsids displayed neurotropic properties in the anterior cingulate cortex, preferentially transducing excitatory neurons (**Figure 5F**), as expected for retrograde capsids following intraputaminal injection. Excitatory projection neurons detected were most likely pyramidal neurons in layer 5 of the cortex, given their characterized projections to the putamen. EGFP mRNA expression was minimally detected in non-neuronal cells of the anterior cingulate (**Figure 5F**), confirming that the vector reached neuronal cell bodies in this distal cortical region via axonal transport from the injection site. The barcode-enriched snRNAseq data were in concordance with RNA amplicon sequencing data for MNM004 and AAV2-retro as the top capsids transducing the anterior cingulate cortex (**Figure 5E**). Our histological data corroborated our bulk RNA barcode amplicon sequencing and snRNA amplicon sequencing data. Coronal brain sections (**Figure 6A**) demonstrated robust EGFP expression in layer 5 neurons of key cortical areas, including anterior cingulate cortex, dorsal prefrontal cortex, and dorsal premotor cortex, consistent with retrograde AAV transport from the putamen.

## DISCUSSION

Neurodegenerative diseases are a group of heterogeneous disorders that mainly affect the CNS. They are characterized by the loss of selectively vulnerable populations of neurons in the brain and spinal cord, causing progressive and incapacitating cognitive, behavioral, and motor dysfunction. Recent advances in neuroimaging techniques have helped delineate the structural and functional connectivity of the human brain and indicated that neurodegenerative diseases tend to target specific brain networks, damaging their function.^7–10^ rAAV gene therapy has shown significant promise in treating various genetic and acquired CNS disorders, for most of which there is currently no cure or effective treatment.^23–25^ One of the most critical challenges in CNS gene therapy development is the precise and efficient targeted delivery of rAAVs to disease-affected brain structures and associated dysfunctional neural networks. Direct IPa delivery into the brain can be used to take advantage of the existing axonal tracts for distant widespread transgene expression in projection neurons of target brain regions.^4,5,26^ However, many natural AAV serotypes do not readily enter axon terminals and therefore have a limited capacity for retrograde transport. Hence, there is a strong need for novel AAV variants with robust retrograde transduction properties in diverse neuronal populations in the brain.^27^ Recent innovations in AAV capsid engineering methods, including directed evolution, rational design, in-silico design, and machine learning, have enabled the discovery of novel CNS capsids.^3,23,28–30^ These AAV capsids are often tested as libraries in *in-vivo* models, such as rodents and NHPs. Because of potential species-specific differences, the challenge with screening designer AAVs is identifying variants with translatable tropism to the human brain. To improve the translatability of novel capsids, testing in NHPs is preferred, as they are animal models most closely related to humans.^12^

In this study, we screened a barcoded library comprised of 29 AAV capsids in the brains of 3 adult cynomolgus monkeys. The library contained 25 AAV2-derived MNM001-MNM025 capsid variants developed using the BRAVE approach, which combines the extensive screening capabilities of directed evolution with the accuracy and reproducibility of rational protein design. The MNM001-MNM025 variants were engineered to display peptides derived from proteins with known affinity for synapses.^11^ These peptides were inserted between amino acids 587 and 588 of the VP(1-3) capsid proteins, thereby disrupting the site of AAV2 binding to HSPGs.^15^ We compared the retrograde spread efficiency, expression levels, and cell-type-tropism of the 25 AAV2-derived variants side-by-side with parental AAV2 and benchmark capsids AAV2-retro, AAV9, and AAV9-retro. The AAV capsid library was bilaterally infused into the putamen, which is a part of the striatum and plays a crucial role in regulating motor functions and cognitive processes.^31^ The extensive and well-defined input and output connections of the putamen with many cortical and subcortical brain areas make it ideally suited to assess AAV retrograde transport properties following IPa delivery. To ensure the specificity and efficacy of our targeting study, it was essential to achieve both sufficient coverage of the putamen and precision of the AAV library delivery, while avoiding vector spillover to adjacent brain structures. We leveraged recent advances in neurosurgical approaches for IPa gene transfer^6^, including convection-enhanced delivery (CED) of MRI contrast-spiked infusate using the reflux-resistant SmartFlow® Neuro Cannula, to monitor and verify AAV vector infusion and target coverage in real time. The AAV vectors and neurosurgical delivery method were well tolerated by the dosed animals, with no neurological deficits or adverse effects noted for the duration of the study. These outcomes are in-line with previously published studies in preclinical large animal models and clinical trials, which assessed the impacts of therapeutic IPa AAV-based gene transfer and reported generally tolerable safety profiles.^4–6,32^ Success in the field was recently marked by the FDA approval of Kebilidi (eladocagene exuparvovec),^33^ as the first gene therapy in the U.S. administered directly into the brain via bilateral intraputaminal infusion for aromatic L-amino acid decarboxylase (AADC) deficiency.^34,35^

The key factor determining the success of gene therapy in patients is the ability to effectively deliver genetic cargo to target brain regions and cell populations. To identify promising novel AAV variants for further development, it is therefore important to accurately characterize their biodistribution and expression level in the NHP brain. Rapid progress in NGS technologies and their significant impact on transcriptomics offered unparalleled capabilities to quantitate and analyze DNA and RNA molecules isolated from cells and tissues.^36^ Our study involved incorporating unique barcodes into each AAV variant within the library, which allowed for high-throughput identification of the most efficient retrograde capsids using bulk RNA barcode amplicon sequencing. The readout of barcodes from RNA ensured that we assessed functional AAV transduction, which encompasses the entire process from viral cell entry to the expression of the transgene. Using this approach, we established MNM004 and AAV2-retro as the two leading retrograde capsids, abundantly expressed throughout the cortical regions and in subcortical brain structures that send projections to the putamen. Our results for MNM004 in the NHP brain agree with previously published findings for MNM004 in the rat brain, where this AAV2 variant displayed robustly improved retrograde neuronal transport relative to parental AAV2.^11^ Cumulatively, these data suggest that axonal properties of MNM004 are conserved across species.

We demonstrated that MNM004 and AAV2-retro displayed similar RNA expression profiles, emerging as the two top-performing retrograde capsids in the NHP brain. These data are consistent with previous observations in the rodent brain.^11^ Interestingly, we discovered that MNM004 retained some expression throughout the putamen, indicating local spread from the injection site. In contrast, AAV2-retro expression in the putamen was essentially absent. We also discovered that AAV2-retro capsid genomes were substantially more abundant in most cortical areas analyzed, compared to MNM004 capsid genomes. These results suggest potential differences between MNM004 and AAV2-retro in their post-cell-entry processing leading to functional transduction. We speculate that the mechanisms involved may include differences in binding to AAV receptor (AAVR), nuclear entry, the rate of AAV uncoating - to release the genome from the capsid, and the efficiency of second strand synthesis - processes previously documented to be both capsid-dependent and rate-limiting in AAV transduction.^37–41^ Additional studies, which are beyond the scope of this manuscript, would be of interest to explore these potential mechanistic differences.

Since the original report by Tervo et al.^13^ that AAV2-retro, which they developed using an *in-vivo* directed evolution approach in mice, demonstrates markedly increased retrograde functionality compared to its parental serotype AAV2, this designer capsid variant was systematically evaluated in several rodent and NHP brain studies.^11,42–44^ Our AAV2-retro data recapitulate published evidence of its strong and extensive axonal retrograde transport to NHP brain areas with direct projections to the striatum.^42–44^ Our data also recapitulate the findings of AAV2-retro not being transported to the occipital lobe and the cerebellum following striatal injection.^44^ In fact, we used cerebellar brain punches as negative experimental controls, as transgene-derived DNA and RNA isolated from these samples did not amplify during sequencing library generation. More recently, Lin et al.^14^ developed an AAV9-retro variant by introducing the 10-mer peptide fragment (LADQDYTKTA) from AAV2-retro into the AAV9 capsid. The authors evaluated AAV9-retro biodistribution in mouse brain after IPa injection and found that it can retrogradely transduce projection neurons with an efficiency comparable to that of AAV2-retro, when these vectors were co-administered into the ventral tegmental area (VTA) or into the striatum.^14^ On the contrary, we showed that in the NHP brain AAV9-retro displayed substantially weaker retrograde transport efficiency compared to that of AAV2-retro. These contrasting findings suggest species-specific differences in receptor/co-receptor for AAV particle uptake at the axon terminals. Finally, our AAV9 expression data are consistent with observations reported by Green et al.^45^ and by Masamizu et al.^46^ in NHPs, which were striatally infused with AAV9 and histologically assessed three or four weeks later, respectively. We found AAV9 to be expressed both locally at the injection site, as well as in cortical and subcortical structures with known projections to the striatum. Historically, AAV9 has been widely used for delivering genetic payloads to the brain due to its efficient neuronal transduction, retrograde and anterograde transport properties, and ability to cross the blood-brain barrier.^47^ In our study, the expression levels of AAV9 at the injection site, cortical regions, and subcortical structures were not superior to those of other capsids in the AAV library. It was notably different from AAV9-retro, however, which was nearly absent from the injection site and from most subcortical structures profiled.

Cell type-specific expression is essential for CNS-directed gene therapies to maximize their efficacy while minimizing off-target effects.^48^ Because our barcoded AAV capsid library is comprised of variants all expressing EGFP transgene driven by the ubiquitous CAG promoter, histological tissue assessment for EGFP protein expression would not reveal cell-type specificity of each capsid variant. We therefore relied on single-nucleus RNA sequencing (snRNAseq) and amplicon enrichment of the unique barcodes to profile AAV tropism. An advantage of the BRAVE-barcoded AAV capsid library is that it can be used without modification in a single-cell/single-nucleus RNA sequencing (sc/snRNAseq) platform,^11^ such as 10X Genomics, where AAV-derived barcoded RNA, driven by a Pol II promoter, is captured via its poly(A) tail. The snRNAseq assay is the preferred way to profile brain tissues, as neurons are fragile and difficult to dissociate. Importantly, this method also allows for application to fresh-frozen tissues.^49^ Because snRNAseq analyzes mostly nuclear transcripts, which represent only a fraction of transcripts in a single cell, the recovery of AAV vector barcodes can become a challenge.^50^ In our study, we enhanced transgene-positive nuclear detection with amplicon enrichment of single-nucleus-encoded EGFP cDNA molecules. This dual approach was applied to selectively profile transduced cell populations of the putamen, substantia nigra, and anterior cingulate cortex. The improvement in the detection of transgene mRNA was especially prominent in the substantia nigra and the anterior cingulate cortex, which had less total AAV-derived transcripts per nucleus compared to the putamen (**Figure S7**).

Dopaminergic neurons in the substantia nigra pars compacta (SNpc) project to the caudate and putamen and form the nigrostriatal pathway, which is crucial for motor control and movement.^31^ Their degeneration is a hallmark of Parkinson’s disease.^51^ AAV-based gene therapies aimed to restore or enhance dopamine signalling in the striatum and the substantia nigra have been actively investigated.^52,53^ Targeting the SNpc for therapeutic gene delivery to nigrostriatal neurons can be risky due to its location deep within the midbrain and its proximity to critical brainstem structures. On the other hand, targeting the striatum is a more feasible surgical approach.^53^ Moreover, a single injection into the putamen allows for therapeutic gene delivery to the nigrostriatal network and thus for a potentially more impactful therapeutic outcome. Therefore, there has been considerable interest in screening both naturally occurring AAV serotypes and designer AAVs for capsid variants with enhanced retrograde properties and preferential transduction of dopaminergic neurons. Several anatomical studies in NHPs characterized AAV targeting to the substantia nigra after striatal delivery.^42–46,54^ These studies, highlighted for their use of the same AAV serotypes as those evaluated as a part of our capsid library, relied on histological brain assessment for transgene protein expression following the infusion of a single AAV into the caudate and/or putamen. It was shown that AAV2 has no retrograde transport properties and instead undergoes anterograde axonal transport from the putamen to the substantia nigra pars reticulata (SNpr).^44,54^ The AAV2-retro-based NHP studies reported varying findings for its retrograde trafficking to the SNpc. Weiss et al.^44^ observed a number of AAV2-retro-positive neurons in the SNpc, although the cell type was not verified by staining for TH, a marker of dopaminergic neurons. In comparison, Albaugh et al.^42^ detected only sparse neuronal labeling, and Cushnie et al.^43^ found no labelled neurons. Finally, AAV9 was demonstrated to undergo bidirectional axonal transport: anterograde, as evidenced by labeling of axon terminals in the SNpr, and retrograde, as evidenced by labeling of neuronal cell bodies in the SNpc.^45,46^

Our bulk RNA amplicon sequencing data did not reveal a specific AAV capsid variant with superior expression in the SN. Because the bulk barcode sequencing approach also had no resolution regarding the cell types transduced, we relied on a barcode enrichment-based snRNAseq strategy to investigate AAV targeting to dopaminergic neurons. Previous studies in rats demonstrated significant retrograde capacity of MNM008 capsid following striatal injection, transducing the great majority of dopaminergic neurons in SNpc.^11^ On the contrary, we found that MNM008 did not retain its favorable characteristics in the NHP brain. The MNM008 capsid was generated by inserting the peptide FTSPLHKNENTV from the CAV-2 capsid. The CAV-2 uses the Coxsackievirus and Adenovirus Receptor (CAR) for infectivity and axonal transport.^55,56^ While the binding capacity of MNM008 to CAR has not yet been determined, it is reasonable to expect that MNM008 would also require the axonal expression and function of CAR, as its retrograde function is derived from the CAV-2 peptide. However, the CAR expression pattern appears to differ between rats and prosimians, such as the grey mouse lemur,^57^ and monkeys, such as the *Macaca fascicularis*.^58^ The CAV-2 capsids have been shown to have significant retrograde transport to the SN in grey mouse lemurs, transducing around 70% of the dopaminergic neurons.^57^ When the same CAV-2 capsids were used in macaques, however, only 5% of the dopaminergic neurons were transduced using the same injection parameters.^58^ With direct injection into the SN, the cells were nonetheless transduced, thus indicating that the axonal expression of the receptor is not as prominent in these larger monkeys. Additionally, we have recently used the MNM008 capsid to efficiently transduce human dopamine neurons in an *in vivo* rat model, allowing the transport from axon terminals in the striatum and prefrontal cortex back into the dopamine transplant in the SN. This indicates that, at least in stem cell-derived human dopamine neurons, the CAR receptor is sufficiently expressed at the axon terminals.^59,60^ Overall, the field needs a deeper understanding of how AAV capsid variants interact with host cells, specifically focusing on receptor and co-receptor usage across different species, brain regions, and neuronal circuits. This will be crucial for optimizing AAV-based gene therapies and understanding viral tropism in the nervous system.

In our study, we established MNM021 as the top capsid transducing dopaminergic neurons in the NHP SN. A limitation of the study is the low number of neuronal nuclei isolated from SN samples, although the transcripts recovered are still highly relevant. Therefore, the results for MNM021 in the SN should be corroborated by a modified snRNAseq screen, where the nuclear population is enriched for neurons using an antibody to NeuN (pan-neuronal sorting) or an antibody to Nurr1 (dopaminergic neuronal sorting).^61^ This modified approach would enrich neuronal nuclei for snRNA-seq characterization, without confounding effects of sequencing the oligodendrocyte population, which is significantly more abundant in the SN.^22^ The MNM021 variant also emerged as the capsid both widely distributed throughout the brain and enriched at the injection site, making it potentially amenable to use in gene therapy applications where it is beneficial to treat both the disease-vulnerable brain region and the associated dysfunctional networks. This approach could enhance the therapeutic benefits in human patients afflicted with Huntington’s disease, Parkinson’s disease, Alzheimer’s disease, or other CNS diseases that affect multiple brain regions and/or neural networks. Specifically with respect to Huntington’s disease, a single injection into the putamen, or into both caudate and putamen, could simultaneously treat the disease’s core pathology, which involves progressive degeneration of the basal ganglia, and the cortico-striatal network, characterized by widespread projections from numerous areas of the cortex to the caudate and putamen. To this end, utilizing retrograde AAVs for single-injection treatments would be more feasible and efficient than administering multiple injections into different cortical areas. This IPa strategy would also facilitate reaching neurons in the deep layers of the cortex, as opposed to injecting AAV vectors into the cerebrospinal fluid (CSF), which often leads to transduction primarily in superficial cortical layers, because the diffusion of AAVs into deeper brain parenchyma is limited by CSF circulation and removal. Importantly, Pressl et al. recently demonstrated that layer 5a corticostriatal pyramidal neurons, which project from the cortex to the striatum, are particularly vulnerable in Huntington’s disease.^62^ They exhibit extensive expansions of the mutant huntingtin (mHTT) gene’s CAG repeats and are selectively lost early in the disease progression, thus causing a disruption of the normal circuit function and potentially contributing to clinical features, such as changes in motor function and cognitive deficits.

Finally, our barcode enrichment-based snRNAseq strategy to profile AAV glial tropism at the injection site yielded inconclusive results. MNM021 was the top capsid variant transducing glial cells in the NHP putamen; however, only in one out of three animals. Therefore, further studies using MNM021 would be of interest to confirm or refute its glial tropism. Notably, MNM017 was previously shown to display a high level of transduction in primary rat glia *in vivo* and in primary human glia *in vitro*.^11^ However, MNM017 did not result in superior transduction of NHP putamen glia, when compared to other capsids in the library. These results suggest species-specific differences in cell surface receptors/co-receptors for AAV that can impact glial cell transduction efficiency.

In summary, we performed a thorough screening of 25 AAV2-based MNM001-MNM025 capsid variants alongside the parental AAV2, AAV2-retro, AAV9, and AAV9-retro capsids in brains of 3 adult cynomolgus monkeys, using DNA- and RNA-derived barcode sequencing, barcode enrichment-based snRNA sequencing and histological assessment. The results of this study demonstrate not only the overall remarkable animal-to-animal consistency, but also assay-to-assay consistency. For instance, MNM004 and AAV2-retro were established in the bulk RNA amplicon sequencing screen as the two top retrograde capsids with superior expression in the anterior cingulate cortex, one of the main brain cortical regions with projections to the striatum.^31^ These results were corroborated by the snRNAseq screen of the anterior cingulate cortex, where MNM004 and AAV2-retro were the two top capsids expressed in the excitatory neurons, which are layer 5 pyramidal neurons, projecting to various cortical and subcortical areas, including the putamen.^31^ Finally, the histological assessment showed an abundance of EGFP-positive neuronal cell bodies in layer 5 of the anterior cingulate cortex, confirming the efficacy and selectivity of the retrograde AAV transport. The results of this study also identified MNM021 as a capsid capable of retrograde trafficking to multiple brain structures, while retaining robust expression at the injection site in the putamen. Further *in vivo* studies exploring this dual functionality of MNM021 in the NHP brain will be of interest to investigate its potential therapeutic relevance and clinical translatability.

## MATERIALS AND METHODS

### AAV library capsid and vector plasmid cloning and amplification

The AAV2-WT backbone plasmid (Spark Therapeutics) was used for the insertion of polypeptide sequences representing MNM001-MNM025 variants into the AAV2-Cap gene at position N587. A separate AAV genome plasmid CAG-EGFP (Spark Therapeutics, Inc.) was used for the insertion of a unique 20 bp molecular barcode downstream of the EGFP reporter gene (**Figure 1A**). Barcoded MNM capsid and AAV genome plasmids were submitted to Aldevron (Fargo, ND) to produce research-grade endotoxin-free giga-scale preps. All plasmids were sequenced-verified by Plasmidsaurus (Eugene, OR) and/or Azenta (Burlington, MA).

### AAV capsid library production

Barcoded recombinant AAVs (rAAVs) were manufactured in the Research Vector Core at Spark Therapeutics. Adherent HEK293 cells were transfected in roller bottles (850 cm^2^) using a standard calcium-phosphate-based triple plasmid system. Each DNA combination contained pHelper, pRepCap, and pTransgene (containing CAG-EGFP and its corresponding unique internal barcode) in a 1:1:1 molar ratio. Twenty-seven vector groups (AAV2 parental, AAV2-retro plus MNM001-MNM025) were transfected individually, because packaging AAV library as a pool for multiplex screening has the risk of barcode recombination.^50,63^ Cells were harvested 96 hours post-transfection. Cell harvests were pooled and processed in groups of 5 or 6 AAVs, i.e. the “mini-libraries.” Cell pellets were collected, resuspended, and lysed by sonication. Clarified cell lysates were digested with benzonase, precipitated with polyethylene glycol (PEG), and purified via 2 rounds of cesium chloride (CsCl) density-gradient ultracentrifugation. Recovered vectors were exchanged to PBS180/0.001% Pluronic F-68 (Gibco, catalog# 24040-032) via cassette dialysis (ThermoFisher, catalog# 66455, 10K MWCO). Vector genomes were quantified by TaqMan qPCR with BGH polyA-specific primers and probe. AAV vectors were characterized by SDS-PAGE/Sypro Ruby stain. The endotoxin level of each vector was less than 1 EU/mL (reported values were < 0.2 EU/mL), measured using Endosafe LAL cartridges and the Endosafe NexGen PTS device (Charles River, Charleston, SC, USA). Vectors were stored at −80°C until use.

AAV9 and AAV9-retro vectors were produced individually and stored at −80°C until use. To produce a complete AAV capsid library for dosing into NHP brain (**Table S1**), the 5 mini-libraries (27 capsids total) and the 2 individually produced capsids were mixed in equal vg amounts for each capsid, for a total of 29 AAVs. An aliquot of the complete AAV capsid library was submitted for next generation sequencing (NGS).

### AAV capsid library characterization

The input capsid libraries were characterized via a UMI-inclusive (quantitative) barcode amplicon sequencing strategy. Briefly, an annealing/extension reaction was performed using a primer with an R2 Illumina sequencing overhang, containing a unique molecular identifier (UMI) and a template-specific sequence (3’ to the barcode). PCR1 was then performed on these products with a primer against the R2 sequencing overhang (which was incorporated in the previous reaction) plus a primer including a R1 sequencing overhang, containing a template-specific sequence 5’ to the barcode. Following this step, indexing PCR was performed, utilizing the now incorporated full-length R1 and R2 sequencing primer binding sites in the molecule. After sequencing, reads were filtered based on the appropriate flanking sequences and then deduplicated via the UMI. Counts were generated for each known barcode to assess relative input capsid abundance.

### Animal welfare statement

The NHP study was conducted at Northern Biomedical Research (NBR) (Norton Shores, MI). All animals were housed and provided enrichment as per NBR’s standard operating procedures. All experimental procedures were reviewed and approved by the NBR Institutional Animal Care and Use Committee. NBR maintains an Animal Welfare Assurance with OLAW and is fully accredited by AAALAC. This study was carried out in accordance with the recommendations in the Guide for the Care and Use of Laboratory Animals of the National Institutes of Health (“The Guide”).

### Serum sample collection

Prior to the initiation of the study, whole blood was collected into serum separator tubes and remained at room temperature until fully clotted. Samples were centrifuged at 2000 rcf at 4°C for 10 min immediately after clotting (within 30 to 45 min of collection). Serum was aliquoted into sterile 1.5 mL safe-lock polypropylene Eppendorf tubes and immediately snap-frozen on dry ice. Aliquoted serum samples were stored in an ultra-low freezer until used in the AAV neutralizing antibody assay.

### AAV neutralizing antibody assay

Neutralizing antibodies (NAbs) against AAV2 and AAV9 were assessed using a cell-based assay with an AAV reporter vector expressing Gaussia luciferase. Serum was heat-inactivated at 56°C for 30 min followed by serial dilution in fetal bovine serum (FBS) to yield a 4-point dilution (1:1 to 1:800). Quality controls (QC) at three levels were prepared using Factor Assay Control (FACT; George King BioMedical, catalog# 0020) in FBS. Positive control (MAX) included reporter vector in DMEM with 50% FBS, while negative control (MIN) contained only DMEM and 50% FBS. The reporter vectors were then added to the diluted QCs, samples, and MAX to achieve a multiplicity of infection (MOI) of either 50 (AAV2) or 350 vg/cell (AAV9). The mixtures were incubated at 37°C for 30 min and subsequently transferred to a 96-well plate containing 2V6.11 cells (20,000 cells/well for AAV2 and AAV9) cultured in DMEM supplemented with 10% FBS, 1X Penicillin-Streptomycin, 1X L-Glutamine, and 1 µg/mL Ponasterone A (ThermoFisher, catalog# H10101). The cells were incubated overnight at 37°C, 5% CO₂, and 20% relative humidity to allow transduction. The following day, 40 µL (AAV2 and AAV9) of supernatant was transferred to a 96-well plate (Corning, catalog# 3915) and analyzed using a GloMax luminometer with the Renilla Luciferase Assay System (Promega, catalog# E2820). Average relative luminescence units (RLU) were calculated for each control and sample. The % luciferase activity was calculated for AAV2 and AAV9: ([Avg sample RLU – Avg MIN RLU/ Avg MAX RLU – Avg MIN RLU]×100). The % inhibition was defined as 100 - % luciferase activity. The NAb titer was reported as the highest dilution with inhibition ≥50%. If a titer could not be assigned due to % inhibition at the last dilution being >50% or the first dilution being <50%, the dilution was reported as >1:dilution or <1:dilution, respectively.

### Surgical delivery of the AAV capsid library

Animals were anesthetized in their home cage, transported to the surgical preparation suite, intubated, and maintained on gas inhalant anesthesia for the duration of the surgical procedure. Viral vector administration was performed using intra-MRI injections targeting the putamen (bilateral) using an occipital trajectory approach. Animals were placed in an MRI-compatible head fixation frame, incisions were made over the calvarium, and craniotomies were created in each hemisphere near the cannula entry points. Animals were transferred to the MRI machine (Philips Achieva 3T MRI) for image collection and dose administration. T1-weighted MRIs were used to identify the target brain regions, and cannula trajectories were planned using Philips software. A small opening in the dura was made by passing a pointed ceramic lancet through the cannula guide to penetrate the dura. A 16-gauge, 10 ft. SmartFlow® Neuro Ventricular Cannula was primed with 150 µL of virus (2x with a 15 min wait period in between). Viral vector was thawed, diluted, and mixed with ProHance (Gadoteridol, 2 mM final concentration) contrast agent to allow for the visualization of vector spread in the putamen throughout the surgical procedure. The cannula was inserted with the infusion pump running at 5 µl/min and lowered to 1 µL/min once at the first infusion site. Vector was delivered at 3 locations along the long axis of the putamen using convection enhanced delivery and a low-ramping infusion rate (1 µL/min for first the 10 min, 2 µL/min for the remainder of the infusion). Animals were injected with a total of 60 µL (animal 1501) or 45 µL (animals 1001 and 1002) per hemisphere and infusions were monitored in real time using a series of short T1W structural scans to visualize Gadoteridol. If vector spread along a vessel was seen during the injection, the cannula was advanced 1-2 mm in attempt to avoid spread of the vector outside the putamen. At the end of the infusion, the cannula had a 5 min dwell time, and upon the cannula removal a final T1W MR image was collected for future quantitation of infusate biodistribution.

### Volumetric analysis of gadoteridol coverage

The percent coverage of Gadoteridol in the putamen was evaluated as a proxy for viral vector distribution throughout the target brain region. T1-weighted structural MRIs acquired pre-infusion, as well as immediately post-infusion, were used to manually segment volumes of the putamen and Gadoteridol using ITK-SNAP software version V 4.0.1.

### Necropsy

Animals were sedated as per contract research organization (CRO) SOP-SRG-001, perfused with chilled PBS (pH 7.4), and biofluids/tissues were collected for molecular and histological studies, including the brain, spinal cord, dorsal root ganglia (DRG), eyes, cerebrospinal fluid (CSF), serum, and plasma.

Following removal from the skull, the brain was placed in a chilled brain matrix and slabbed coronally at approximately 4 mm thickness; 17 slabs total were generated per animal. Each brain was then hemisected into left and right hemisphere. For animals 1001 and 1002, fresh-frozen brain samples were collected from the left hemisphere, whereas the right hemisphere was processed for immunohistochemistry. For animal 1501, fresh-frozen brain samples were collected from both hemispheres.

For fresh-frozen brain sample collection, the coronal brain slabs were laid out in petri dishes and samples were collected according to a pre-determined punch diagram, using 3 mm diameter biopsy punches (EMS, catalog# 69031-45). Selected areas were micro-dissected: substantia nigra, globus pallidus, and cerebellar cortex. All brain samples were collected into 2.0 mL TPP® cryotubes (Millipore Sigma, catalog# Z760951), weighed, snap-frozen in liquid nitrogen, and stored at -80°C until use. A total of 80 samples were collected per brain hemisphere. The entire study included 240 brain samples. The residual brain slabs were placed between 2 pieces of parafilm into foil envelopes and immediately frozen by completely submerging in dry ice. Upon freezing, the brain slabs were transferred to -80°C for storage.

For histology, the coronal slabs from the right hemisphere were post-fixed in 10% neutral buffered formalin (NBF) at room temperature for 24 ± 4 hours with gentle shaking, then transferred to 70% ethanol and saved refrigerated until processed for formalin-fixed paraffin-embedding (FFPE). The embedding occurred within 10 days of tissue collection. All tissue processing for both molecular and histological studies took place within a 2-hour post-mortem interval.

### Tissue processing for DNA and RNA

Fresh-frozen brain samples were processed for dual DNA/RNA extraction using miRNeasy Advanced Mini Kit (Qiagen, catalog# 217604) as per manufacturer’s instructions. Briefly, tissues were homogenized in lysis buffer using Tissuelyser II (Qiagen, catalog# 85300) and 5 mm stainless steel beads (Qiagen, catalog# 69989). Lysate was transferred to a genomic DNA (gDNA) eliminator column, and the flow-through was collected and processed for total RNA extraction. RNA was treated off-column with TURBO DNAse (ThermoFisher, catalog# AM1907) to eliminate residual genomic and vector-derived DNA, aliquoted and stored at -80°C until use. As a modification to the manufacturer’s protocol, the gDNA eliminator columns were saved and DNA was extracted using a standard procedure based on Proteinase K (Qiagen, catalog# 19131) and buffers AW1 and AW2 (Qiagen, catalog# 19081 and 19072, respectively). Extracted DNA was aliquoted and stored at -20°C until use. RNA and DNA concentration and purity were assayed using the NanoDrop 2000 (ThermoFisher, catalog# ND-2000). RNA integrity number (RIN) values were obtained using the 4200 Agilent TapeStation (Agilent, catalog# G2991BA).

### Tissue processing for snRNAseq

For each brain structure analyzed, 2 fresh-frozen tissue punches (putamen, anterior cingulate cortex) and 2 fresh-frozen micro-dissected areas (substantia nigra) from adjacent coronal slabs were pooled into a single sample with a combined weight of ≤ 50 mg. For each brain structure, samples were analyzed from a minimum of two animals. Nuclei were isolated using the 10x Genomics Nuclei Isolation Kit with RNase Inhibitor (10x Genomics, catalog# 1000494) with minor modifications. First, flash-frozen tissue was transferred to ice-cold lysis buffer (from the Nuclei Isolation Kit) in a 1 mL glass tissue homogenizer (Wheaton, catalog# 357538). After dounce homogenization, the tissue lysate was resuspended and left to incubate for 10 minutes on ice. Additional steps followed manufacturer’s instructions. During the first wash (after debris removal), the nuclear suspension was incubated with rotation for 20 min at 4°C with DAPI staining solution (Abcam, catalog# ab228549) at a final concentration of 5 µM. Following a spin and final resuspension, nuclei with 2N DNA content were purified by fluorescence-activated nuclear sorting (Sony Cell Sorter) based on DAPI fluorescence and collected in Wash and Resuspension Buffer (PBS, 0.5% BSA, and 0.2U/µL RNase Inhibitor - prepared according to kit instructions). Sorted 2N nuclei event count and final volume were used to inform sample loading volume, and the nuclei were immediately loaded onto 10x Chromium Single Cell 3’ Kit v3.1 chips (10x Genomics, catalog# 1000268) at ∼20k sorter nuclear event counts per sample (this loading concentration was optimized based on sorted event count and 10x nuclear recovery). Libraries were generated following the manufacturer’s protocol and were sequenced using the Illumina NextSeq 2000.

Dzhashiashvili et al.

### AAV capsid biodistribution analysis

#### Bulk DNA and RNA Barcode Characterization

Bulk DNA and RNA barcode counts were generated using the same quantitative (UMI-inclusive) sequencing strategy as for the input vector libraries. For DNA, bulk total DNA was input into the R2+UMI annealing/extension reaction. For RNA, bulk total RNA was input to a reverse transcription (RT) reaction where our RT primer included the R2 sequencing overhang + a UMI + an anchored poly(dT) sequence. Following this step, we used endpoint qPCR to read out relative template abundance (using the same PCR1 primers as in the vector characterization experiment). Ct values from this were used to inform total amplification cycles for the library preparation strategy (which are split between PCR1 and indexing PCR). The same PCR1 and indexing PCR conditions were used for bulk RNA/DNA libraries as for the input vector libraries. For bioinformatics analysis, UMIs were extracted and reads filtered by flanking sequences to make sure the amplicons were valid. Barcode sequences were extracted according to fixed flanking sequences and reads with the same UMI and barcode sequences were collapsed as PCR duplicates. De-duplicated counts were obtained for each BRAVE barcode.

#### snRNAseq + snRNA Barcode Characterization

We used the 10x Genomics v3.1 3’ GEX kit to generate full-length, barcoded cDNA from nuclei isolated from select brain regions of interest. After generating full-length cDNA, 25% of the cDNA underwent the traditional whole-transcriptome library preparation protocol, where the remaining cDNA underwent a BRAVE barcode amplicon enrichment strategy. Briefly, PCR1 was performed with a template-specific primer + an R2 sequencing overhang as well as a primer against the R1 sequencing overhang that was added during the 10x RT barcoded priming step. The resulting product includes the R1 sequencing overhang, UMI, 10x cellular barcode, BRAVE barcode (at a fixed position within the fragment), and R2 overhang. This combination enriches for BRAVE barcode information that can be attributed to specific cell types using matched 10x barcode information between this amplicon dataset and the whole-transcriptome snRNA-seq data, since both datasets were generated from the same amplified, barcoded cDNA pool. For bioinformatics analysis, the snRNA reads were processed with CellRanger (v7.0); clustering and cell type annotations were performed in Seurat (v4.9). The barcode enrichment amplicon reads were processed similarly as the bulk characterization detailed above. The snRNA and amplicon enrichment datasets were connected via filtered 10x cellular barcodes present in each dataset. Non-cellular background was used to filter BRAVE barcode counts in true cells, where for each BRAVE barcode in each true cell, only barcode counts higher than the 98% quantile of the same barcode in non-cellular units were retained.

### Histology and immunohistochemistry

Coronal brain slabs were processed for FFPE by the CRO and shipped to Spark Therapeutics as paraffin blocks. Paraffin-embedded brain slabs were cut into 5 µm sections onto Fisherbrand Superfrost Plus slides (Fisher, catalog# 12-550-15) and immunostained for green fluorescent protein (GFP), nuclear neuronal protein (NeuN), and tyrosine hydroxylase (TH). The immunohistochemistry protocol was performed as follows. Sections were baked in the ACD HybEZ™ II Oven (Biotechne, catalog# 321710) for 1 hour at 60°C to melt the paraffin, deparaffinized in Histo-Clear (Fisher Scientific, catalog# 50-899-90147), rehydrated through a 100%, 95%, 70%, and 50% ethanol series, and rinsed in distilled water. Heat-induced epitope retrieval was performed using pH 6 citrate buffer (BioSB, catalog# BSB 0020) in a TintoRetriever Digital Pressure Cooker (BioSB, catalog# BSB 7008) at 100°C with low pressure setting for 20 minutes. Subsequently, the slides were cooled for 30 min at room temperature and washed three times in 1X PBS, 5 minutes each. Tissue sections were then incubated in blocking solution (5% BSA, 1% donkey serum, and 0.2% Triton X-100 in 1X PBS) for 1 hour at room temperature. Primary antibodies to GFP (1:2,000; Abcam, catalog# ab6673), NeuN (1:500; Abcam, catalog# ab104224), and TH (1:500; Pel-Freez biologicals, catalog# P40101-150) were diluted in the blocking solution, and tissue sections were incubated overnight at 4°C. The next day, the sections were washed once in 1X PBS containing 0.2% Triton X-100 and then twice in 1X PBS, for a total of three washes, 10 minutes each. Secondary antibodies were applied in the blocking solution for 2 hrs at room temperature. The following secondary antibodies were used: donkey anti-goat 488 (1:500; ThermoFisher, catalog# A32814), donkey anti-rabbit 555 (1:300; ThermoFisher, catalog# A32794), and donkey anti-mouse 647 (1:300; ThermoFisher, catalog# A32787). After removing the secondary antibodies, the sections were washed in 1X PBS containing 0.2% Triton X-100 followed by washes in 1X PBS as described above, and autofluorescence blocking was performed for 3 minutes using Vector® TrueVIEW® Autofluorescence Quenching Kit (Vector Labs, catalog# SP-8400-15) as per manufacturer’s instructions. Immediately after, the sections were rinsed in distilled water and mounted in Prolong Gold medium with 4’,6-diamidino-2-phenylindole (DAPI) (ThermoFisher, catalog# P36931). Slides with sections were imaged on a digital slide scanner (Zeiss Axioscan Z7) at 10X magnification, and images were transferred to the HALO viewing platform.

## DATA AND CODE AVAILABILITY

All data and code are available upon a reasonable request to the corresponding authors.

## ACKNOWLEDGEMENTS

This work was funded by Spark Therapeutics, a subsidiary of Hoffman-La Roche. The authors thank Spark Therapeutics Research Vector Core (Xin Zhang, Lihua Shi, and Charmaine Azutillo), Histology Core (Ashley Carter and Renee Gentzel), Computational Biology and Machine Learning (Perry Evans), Bioanalytical Sciences (Mallory Becker and Heena Beck), and Scientific Writing (Aurore Lebrun) for experimental, data analysis, and data visualization support.

## AUTHOR CONTRIBUTIONS

Conceptualization, T.B., M.D., and E.A.E.; funding acquisition, E.A.E. and E.R.; methodology, Y.D., J.L.M., E.F., X.H., Z.Y., P.R. and M.N.; investigation, Y.D., J.L.M., E.F., X.H., B.M.K., G.D.W., M.N. and A.A.H.; visualization, Y.D., J.L.M., E.F., X.H., A.A.H. and M.N.; project administration and supervision, Y.D. and E.A.E.; software, X.H., A.A.H., and P.R.; validation, Y.D., J.L.M., E.F., X.H. and A.A.H.; writing – original draft, Y.D., J.L.M., E.F., X.H. and T.B.; writing –review and editing, Y.D., J.L.M., E.F., X.H., B.M.K., G.D.W., M.N., A.A.H., Z.Y., P.R., E.R., M.D., E.A.E., and T.B.

