## Supplemental Tables & Figures for "Administration of barcoded AAV capsid library to the putamen of non-human primates identifies variants with efficient retrograde transport"

### SUPPLEMENTAL INFORMATION

#### Table S1. Barcoded AAV capsid library

| AAV | Serotype |
|---|---|
| Capsid (AAV2-based) | MNM001 – MNM025 |
| Capsid (benchmark) | AAV2 (parental), AAV2-retro, AAV9, AAV9-retro |

#### Table S2. Peptide identity of AAV2-derived BRAVE capsid library.

| Capsid | Family | Protein | Name | Ref seq |
|---|---|---|---|---|
| MNM001 | Herpes simplex virus type 2 | Envelope glycoprotein C | HSV-2-pUL44 | JN561323.2 |
| MNM002 | Herpes simplex virus type 2 | Envelope glycoprotein L | HSV-2-pUL1 | JN561323.2 |
| MNM003 | Herpes simplex virus type 2 | Envelope glycoprotein B | HSV-2-pUL27 | JN561323.2 |
| MNM004 | Herpes simplex virus type 2 | Envelope glycoprotein H | HSV-2-pUL22 | JN561323.2 |
| MNM005 | Enterovirus 71 | Capsid protein | EV71-VP3 | AB239756.1 |
| MNM006 | Herpes simplex virus type 1 | Inner tegument protein | HSV-1-pUL37 | JQ673480.1 |
| MNM007 | Herpes B virus | Glycoprotein D | BV-G | S48101.1 |
| MNM008 | Canine adenovirus 2 | Fiber | CAV-2-F-SH01 | KF727562.1 |
| MNM009 | AD & PD proteins | Microtubule-associated protein Tau | Tau | M84156.1 |
| MNM010 | Neurotoxins | Botulinum neurotoxin | BoNT/E Hc | Q00496.2 |
| MNM011 | Tick-borne encephalitis virus | E protein | TBEV-E4387/B7 | X76608.1 |
| MNM012 | Measles virus | Hemagglutinin | MV-H-Edmonston | AF266288 |
| MNM013 | Horseradish peroxidase | Horseradish peroxidase | HRP | A00740.1 |
| MNM014 | Adeno-associated virus | Capsid protein | AAV1-VP1 | AF063497.1 |
| MNM015 | Pseudorabies virus | Glycoprotein E | PRV-Bercker-gE | JF797219.1 |
| MNM016 | Herpes simplex virus type 1 | Envelope glycoprotein D | HSV-1-pUS6 | JQ673480.1 |
| MNM017 | AD & PD proteins | Microtubule-associated protein Tau | Tau | M84156.1 |
| MNM018 | Herpes simplex virus type 2 | Envelope glycoprotein D | HSV-2-pUS6 | JN561323.2 |
| MNM019 | Canine adenovirus 2 | Fiber | CAV-2-F-SH01 | KF727562.1 |
| MNM020 | Adeno-associated virus | Capsid protein | AAV1-VP1 | AF063497.1 |
| MNM021 | Herpes simplex virus type 2 | Envelope glycoprotein M | HSV-2-pUL10 | JN561323.2 |
| MNM022 | Herpes simplex virus type 2 | Major capsid protein | HSV-2-pUL19 | JN561323.2 |
| MNM023 | Herpes simplex virus type 1 | Envelope glycoprotein H | HSV-1-pUL22 | JQ673480.1 |
| MNM024 | Neurotoxins | Botulinum neurotoxin | BoNT/C Hc | 3R4U_A, BAM65691.1 |
| MNM025 | Lectins | Soybean agglutinin | SA | 1SBF_A |

#### FIGURE LEGENDS

##### Figure S1. The MNM004 capsid promotes highly efficient retrograde transport

Visual representation (heatmaps) of RNA amplicon sequencing data for MNM004, parental AAV2 and benchmark AAV2-retro capsids. For each capsid, data were aggregated from 2 NHPs (for substantia nigra) or from 3 NHPs (for all other brain samples). Diagrams demonstrate samples profiled within consecutive rostral-to-caudal coronal brain slabs. For each brain slab, a corresponding anatomical atlas brain map is displayed for reference. The actual numeric NGS data for each brain sample (% barcode variant abundance) were directly and automatically converted into colored pixels with red indicating higher expression values and blue indicating signal values below the threshold. The intensity of the red/blue palette correlates with the numeric values. Brain samples were collected as 3 mm punches, except globus pallidus and substantia nigra, which were microdissected and either split in half (external and internal globus pallidus) or collected as a whole (substantia nigra). For these three structures, colored circles represent data from dissected tissues. NHP, non-human primate; NGS, next-generation sequencing.

##### Figure S2. RNA and DNA amplicon sequencing data comparing expression and biodistribution of select capsids in NHP brain

Amplicon sequencing data comparing expression (RNA) and biodistribution (DNA) of select capsids in brain cortical regions and subcortical structures. Data for each capsid (MNM004, MNM021, parental AAV2, and benchmark AAV9 and AAV2-retro) are shown side-by-side; data from averaged from 2 NHPs (for substantia nigra) or from 3 NHPs (for all other brain samples). Except for amygdala, 2 or more samples (punches or dissected areas) per brain structure were collected and analyzed. Each row in the RNA and DNA data tables corresponds to individual brain sample data that show percent capsid barcode variant abundance, out of 100% total for the entire library. NHP, non-human primate.

##### Figure S3. RNA amplicon sequencing data comparing expression of select capsids in NHP brain

Amplicon sequencing data reporting expression (RNA) of select capsids in brain cortical regions and subcortical structures. Data for each capsid are shown side-by-side and averaged from 2 NHPs (for substantia nigra) or from 3 NHPs (for all other brain samples). Except for amygdala, 2 or more samples (punches or dissected areas) per brain structure were collected and analyzed. Each row in the RNA data table corresponds to individual brain sample data that show percent capsid barcode variant abundance, out of 100% total for the entire library. NHP, non-human primate.

##### Figure S4. Cell-type-specific genes and cluster annotations for NHP putamen

(A) Dot plot of cell marker genes detailing average expression and percentage of cells expressing each marker gene at the transcript level. The numbers on the Y-axis correspond to clusters in (B) UMAP of cell types identified in the putamen by snRNAseq. MSN1, medium spiny neurons type 1; MSN2, medium spiny neurons type 2; OPC, oligodendrocyte precursor cells; UMAP, Uniform Manifold Approximation and Projection.

##### Figure S5. Cell-type-specific genes and cluster annotations for NHP substantia nigra

(A) Dot plot of cell marker genes detailing average expression and percentage of cells expressing each marker gene at the transcript level. The numbers on the Y-axis correspond to clusters in (B) UMAP of cell types identified in the substantia nigra by snRNAseq. Oligos, oligodendrocytes; OPC, oligodendrocyte precursor cells; PVM, perivascular macrophages; UMAP, Uniform Manifold Approximation and Projection.

##### Figure S6. Cell-type-specific genes and cluster annotations for NHP anterior cingulate cortex

(A) Dot plot of cell marker genes detailing average expression and percentage of cells expressing each marker gene at the transcript level. The numbers on the Y-axis correspond to clusters in (B) UMAP of cell types identified in the anterior cingulate cortex by snRNAseq. Oligos, oligodendrocytes; OPC, oligodendrocyte precursor cells; PVM, perivascular macrophages; Ast, astrocytes; UMAP, Uniform Manifold Approximation and Projection.

##### Figure S7. Relative detection of transgene(+) nuclei via traditional 10x 3’ GEX versus amplicon enrichment sequencing

Bar plots detailing relative detection of transgene(+) nuclei number (left) and percentages (right) via traditional 10x 3’ GEX (blue) versus amplicon enrichment sequencing of the transgene from single-nucleus cDNA (orange) for various cell types in the NHP. (A) Putamen (injection site, N=3). (B) Anterior Cingulate (N=3), and (C) Substantia Nigra (N=2). Percentage in this figure is defined as the number of transgene(+) cells detected via either method within a cell type, divided by the total transgene(+) cells detected in that sample.

#### Figure S1. The MNM004 capsid promotes highly efficient retrograde transport

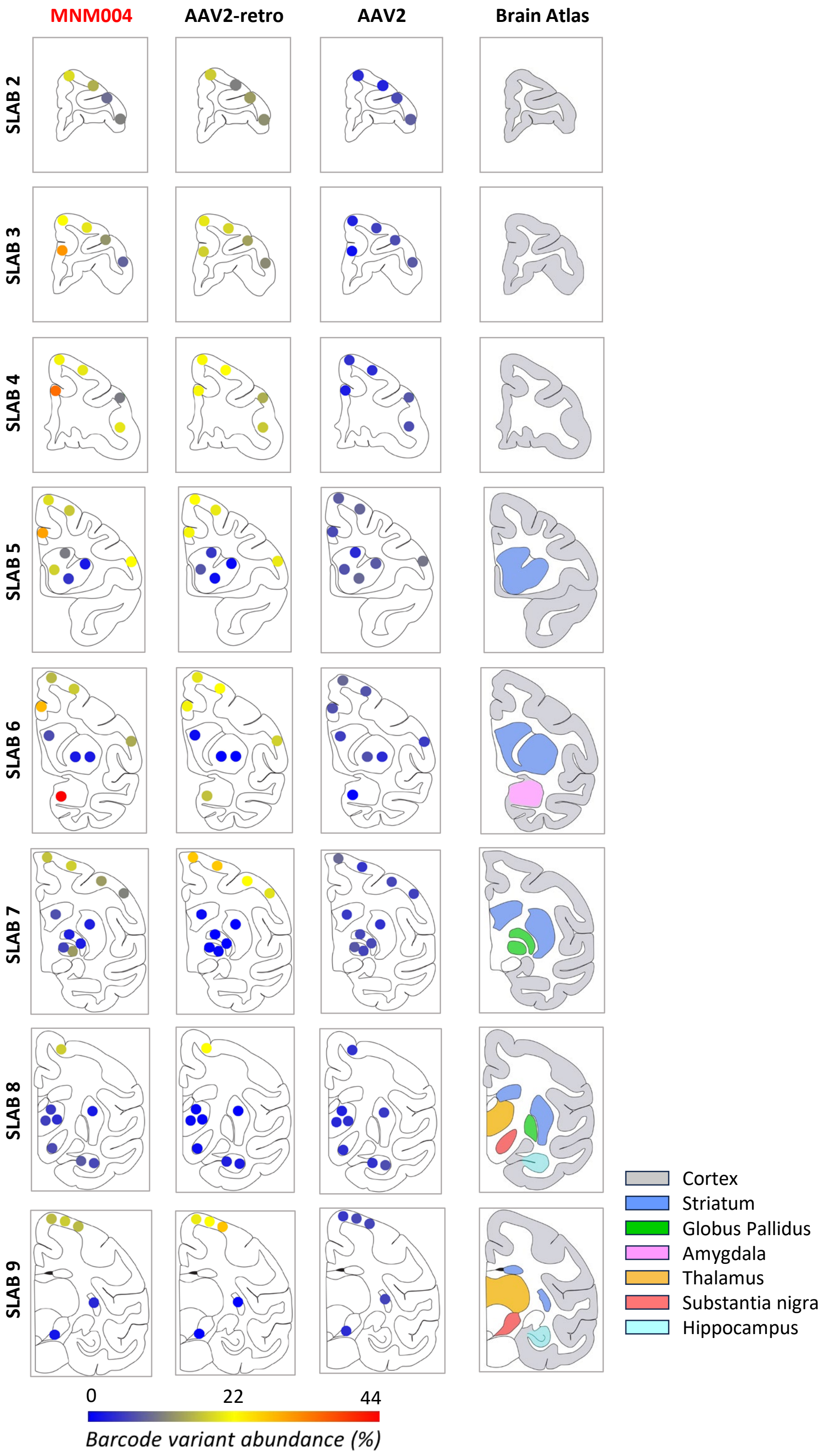

### Figure S2. RNA and DNA amplicon sequencing data comparing expression and biodistribution of select capsids in NHP brain

CORTEX: Dorsal Prefrontal Cortex through Somatosensory Cortex

INJECTION SITE: Putamen

**RNA amplicon seq**

| BRAIN STRUCTURE | AAV2 | AAV2-retro | MNM004 | MNM021 | AAV9 | AAV9-retro |
|---|---|---|---|---|---|---|
| Dorsal Prefrontal Cortex | 0.93 | 16.50 | 17.77 | 6.18 | 3.05 | 0.65 |
| | 0.77 | 10.34 | 14.15 | 5.82 | 3.52 | 1.13 |
| | 0.73 | 18.69 | 20.43 | 5.64 | 2.02 | 1.12 |
| | 1.31 | 17.24 | 18.47 | 6.04 | 1.50 | 1.49 |
| | 1.53 | 13.08 | 11.97 | 8.33 | 3.55 | 1.44 |
| | 1.05 | 20.21 | 19.66 | 5.52 | 2.04 | 1.52 |
| | 0.94 | 20.85 | 18.64 | 5.51 | 2.11 | 1.49 |
| | 1.71 | 13.99 | 10.01 | 6.28 | 2.26 | 1.03 |
| Ventral Prefrontal Cortex | 1.49 | 12.62 | 8.89 | 6.07 | 1.98 | 0.98 |
| | 1.87 | 11.50 | 10.36 | 6.22 | 1.98 | 1.84 |
| | 1.77 | 11.21 | 7.98 | 6.96 | 2.14 | 1.39 |
| | 1.55 | 16.15 | 18.47 | 5.43 | 2.18 | 1.61 |
| Anterior Cingulate Cortex | 0.41 | 16.72 | 28.74 | 4.99 | 1.74 | 0.48 |
| | 0.63 | 19.92 | 32.84 | 4.35 | 1.57 | 1.05 |
| | 1.49 | 19.87 | 27.52 | 4.46 | 1.70 | 1.85 |
| | 1.62 | 19.64 | 25.21 | 5.03 | 1.56 | 1.50 |
| Dorsal Premotor Cortex | 1.83 | 20.76 | 17.92 | 5.17 | 1.91 | 2.47 |
| | 2.00 | 18.82 | 16.12 | 4.64 | 2.34 | 2.13 |
| | 2.05 | 18.47 | 14.85 | 5.30 | 2.17 | 3.87 |
| | 1.67 | 20.85 | 16.29 | 5.43 | 1.59 | 2.48 |
| Ventral Premotor Cortex | 2.28 | 18.94 | 20.53 | 4.92 | 1.85 | 2.29 |
| | 1.21 | 16.81 | 13.87 | 6.18 | 2.86 | 2.03 |
| Primary Motor Cortex | 2.07 | 23.73 | 15.72 | 5.24 | 1.83 | 3.43 |
| | 1.13 | 23.66 | 16.63 | 5.64 | 1.60 | 2.45 |
| | 1.05 | 21.19 | 16.54 | 5.00 | 2.18 | 1.96 |
| | 1.16 | 18.96 | 15.01 | 5.40 | 2.23 | 2.59 |
| | 1.47 | 21.22 | 16.40 | 5.40 | 2.01 | 2.41 |
| | 1.39 | 24.16 | 14.99 | 5.22 | 1.61 | 6.83 |
| Somatosensory Cortex | 1.43 | 20.52 | 12.58 | 5.63 | 2.19 | 4.00 |
| | 1.33 | 18.24 | 10.57 | 7.05 | 2.49 | 2.21 |
| Caudate | 0.91 | 4.58 | 10.02 | 5.40 | 2.11 | 3.84 |
| | 1.63 | 6.92 | 16.80 | 4.12 | 2.38 | 1.99 |
| | 1.27 | 1.08 | 6.27 | 7.86 | 2.49 | 1.78 |
| | 1.17 | 0.95 | 6.52 | 7.91 | 3.03 | 0.86 |
| Putamen | 1.77 | 0.71 | 1.98 | 4.80 | 4.85 | 0.45 |
| | 2.04 | 1.12 | 4.28 | 3.71 | 3.84 | 0.57 |
| | 0.93 | 0.16 | 2.24 | 8.60 | 4.43 | 0.23 |
| | 1.62 | 0.10 | 2.13 | 9.70 | 3.66 | 0.15 |
| | 1.04 | 0.18 | 2.59 | 9.51 | 4.46 | 0.24 |
| | 1.17 | 0.25 | 2.14 | 9.35 | 4.68 | 0.30 |
| | 1.37 | 0.22 | 3.37 | 9.21 | 3.73 | 0.25 |
| External Globus Pallidus | 1.42 | 0.11 | 2.54 | 7.88 | 4.94 | 0.38 |
| | 1.46 | 0.17 | 2.28 | 8.30 | 6.10 | 0.44 |
| Internal Globus Pallidus | 1.74 | 0.29 | 5.49 | 6.57 | 3.74 | 0.28 |
| | 1.38 | 0.68 | 12.63 | 5.19 | 4.13 | 0.35 |
| Amygdala | 0.23 | 15.60 | 43.91 | 3.23 | 1.56 | 0.98 |
| Thalamus | 0.74 | 0.51 | 4.69 | 7.84 | 3.78 | 0.30 |
| | 0.86 | 0.80 | 5.28 | 8.46 | 4.67 | 0.24 |
| | 1.00 | 0.36 | 3.85 | 8.55 | 4.12 | 0.25 |
| Hippocampus | 1.56 | 2.18 | 5.82 | 9.98 | 5.03 | 0.54 |
| | 1.06 | 2.67 | 7.35 | 8.94 | 4.25 | 0.40 |
| Substantia Nigra | 1.46 | 1.35 | 9.24 | 6.78 | 6.15 | 0.70 |
| | 1.43 | 0.14 | 2.70 | 7.31 | 6.42 | 0.24 |

**DNA amplicon seq**

| BRAIN STRUCTURE | AAV2 | AAV2-retro | MNM004 | MNM021 | AAV9 | AAV9-retro |
|---|---|---|---|---|---|---|
| Dorsal Prefrontal Cortex | 0.31 | 1.46 | 0.67 | 1.02 | 6.47 | 0.60 |
| | 0.29 | 0.83 | 0.73 | 2.15 | 6.59 | 0.14 |
| | 0.55 | 2.74 | 1.35 | 2.25 | 6.89 | 0.65 |
| | 1.06 | 2.95 | 1.27 | 1.95 | 7.42 | 1.42 |
| | 0.91 | 7.73 | 2.34 | 2.21 | 6.00 | 2.43 |
| | 0.61 | 3.72 | 1.50 | 1.65 | 7.00 | 0.75 |
| | 1.24 | 2.31 | 0.09 | 1.15 | 4.24 | 0.55 |
| Ventral Prefrontal Cortex | 0.56 | 2.39 | 0.41 | 0.55 | 5.62 | 0.60 |
| | 0.20 | 4.10 | 0.71 | 2.11 | 4.06 | 0.53 |
| | 2.14 | 5.59 | 2.30 | 1.26 | 3.50 | 1.22 |
| | 2.08 | 33.14 | 4.99 | 1.45 | 2.25 | 5.44 |
| Anterior Cingulate Cortex | 0.24 | 1.50 | 1.36 | 1.70 | 5.75 | 0.56 |
| | 0.48 | 5.53 | 2.30 | 1.90 | 6.52 | 0.96 |
| | 1.81 | 22.71 | 5.65 | 1.79 | 4.31 | 6.02 |
| | 2.27 | 23.10 | 4.47 | 2.13 | 4.55 | 3.20 |
| Dorsal Premotor Cortex | 2.04 | 21.32 | 3.11 | 1.79 | 2.74 | 3.98 |
| | 1.66 | 12.44 | 1.99 | 1.46 | 4.95 | 2.44 |
| | 2.22 | 33.31 | 4.42 | 1.60 | 2.13 | 6.45 |
| | 1.97 | 14.03 | 2.41 | 1.54 | 2.77 | 2.63 |
| Ventral Premotor Cortex | 2.47 | 42.83 | 6.28 | 2.14 | 1.67 | 7.13 |
| | 1.39 | 20.63 | 2.88 | 2.42 | 1.83 | 4.12 |
| Primary Motor Cortex | 2.56 | 40.94 | 3.35 | 1.99 | 1.58 | 9.23 |
| | 2.28 | 22.54 | 3.43 | 1.74 | 2.38 | 3.87 |
| | 1.88 | 17.45 | 4.47 | 2.88 | 3.92 | 2.66 |
| | 1.35 | 22.77 | 3.38 | 2.25 | 3.41 | 3.13 |
| | 1.40 | 25.92 | 5.30 | 1.58 | 2.71 | 3.89 |
| | 1.95 | 42.35 | 2.90 | 1.99 | 2.17 | 5.63 |
| Somatosensory Cortex | 2.05 | 28.84 | 2.35 | 2.08 | 3.15 | 7.83 |
| | 1.49 | 15.62 | 4.04 | 2.83 | 3.61 | 3.32 |
| Caudate | 1.16 | 5.05 | 2.45 | 1.63 | 4.43 | 2.62 |
| | 0.65 | 4.82 | 2.69 | 0.81 | 4.91 | 2.86 |
| | 1.56 | 3.47 | 1.91 | 2.90 | 3.84 | 3.15 |
| | 1.16 | 6.24 | 1.86 | 1.53 | 3.73 | 1.66 |
| Putamen | 1.76 | 3.15 | 1.91 | 3.22 | 4.39 | 1.99 |
| | 1.49 | 3.84 | 2.38 | 2.16 | 3.97 | 1.69 |
| | 2.76 | 3.18 | 1.36 | 4.67 | 4.45 | 4.43 |
| | 7.31 | 4.82 | 1.26 | 3.92 | 3.47 | 13.04 |
| | 4.29 | 4.37 | 1.38 | 4.56 | 4.21 | 6.59 |
| | 2.62 | 2.80 | 1.32 | 5.83 | 4.42 | 2.73 |
| | 4.98 | 4.39 | 1.54 | 4.85 | 3.41 | 6.87 |
| External Globus Pallidus | 1.12 | 2.01 | 1.14 | 4.23 | 5.63 | 1.11 |
| | 2.21 | 2.63 | 1.35 | 4.42 | 6.21 | 2.07 |
| Internal Globus Pallidus | 1.24 | 1.11 | 1.38 | 2.49 | 5.49 | 0.51 |
| | 1.02 | 0.48 | 0.98 | 2.43 | 5.81 | 0.44 |
| Amygdala | 0.93 | 7.40 | 8.93 | 1.00 | 3.38 | 1.64 |
| Thalamus | 0.62 | 0.59 | 2.47 | 1.70 | 2.50 | 0.67 |
| | 0.46 | 1.22 | 1.96 | 1.55 | 5.90 | 1.03 |
| | 0.52 | 0.31 | 0.86 | 1.18 | 1.74 | 0.45 |
| Hippocampus | 0.31 | 1.29 | 1.71 | 5.50 | 1.18 | 1.30 |
| | 0.90 | 0.07 | 0.71 | 3.63 | 5.30 | 0.28 |
| Substantia Nigra | 0.95 | 0.04 | 1.09 | 2.28 | 6.88 | 0.56 |
| | 0.88 | 0.03 | 1.00 | 2.52 | 6.90 | 0.40 |

0 — 22 — 44

*Barcode variant abundance (%)*

### Figure S3. RNA amplicon sequencing data comparing expression of select capsids in NHP brain

CORTEX: Dorsal Prefrontal Cortex – Somatosensory Cortex

INJECTION SITE: Putamen

| BRAIN STRUCTURE | AAV2 | MNM001 | MNM002 | MNM003 | MNM005 | MNM006 | MNM007 | MNM008 | MNM009 | MNM010 | MNM011 | MNM012 | MNM013 | MNM014 | MNM015 | MNM016 | MNM017 | MNM018 | MNM019 | MNM020 | MNM021 | MNM022 | MNM023 | MNM024 | MNM025 | AAV9 | AAV9-retro | BRAIN STRUCTURE |
|---|---|---|---|---|---|---|---|---|---|---|---|---|---|---|---|---|---|---|---|---|---|---|---|---|---|---|---|---|
| Dorsal Prefrontal Cortex | 0.93 | 2.36 | 1.27 | 1.92 | 1.81 | 0.92 | 1.52 | 2.00 | 2.08 | 1.10 | 1.69 | 3.73 | 3.46 | 0.97 | 3.62 | 2.04 | 3.22 | 4.57 | 2.50 | 1.42 | 6.18 | 2.26 | 1.10 | 2.42 | 3.99 | 3.05 | 0.65 | Dorsal Prefrontal Cortex |
| | 0.77 | 2.35 | 1.83 | 2.37 | 1.76 | 0.69 | 2.04 | 2.12 | 1.73 | 0.20 | 1.39 | 5.19 | 4.24 | 0.88 | 4.45 | 3.28 | 3.16 | 4.29 | 2.81 | 2.28 | 5.82 | 4.57 | 1.25 | 3.37 | 4.10 | 3.52 | 1.13 | |
| | 0.73 | 1.89 | 1.28 | 1.34 | 1.94 | 0.67 | 1.85 | 1.89 | 2.38 | 0.35 | 1.55 | 3.64 | 4.12 | 0.33 | 3.28 | 2.51 | 3.06 | 3.70 | 1.45 | 1.93 | 5.64 | 2.05 | 1.34 | 3.37 | 2.88 | 2.02 | 1.12 | |
| | 1.31 | 2.08 | 1.48 | 2.17 | 2.10 | 1.33 | 1.85 | 2.03 | 2.51 | 0.73 | 2.01 | 4.07 | 3.54 | 0.93 | 3.27 | 2.07 | 3.08 | 3.91 | 1.42 | 1.82 | 6.04 | 2.34 | 1.27 | 3.29 | 2.91 | 1.50 | 1.49 | |
| | 1.53 | 2.72 | 1.87 | 2.51 | 2.36 | 1.02 | 1.81 | 1.45 | 2.40 | 0.34 | 2.56 | 4.12 | 4.93 | 0.43 | 4.27 | 2.20 | 2.84 | 6.47 | 0.94 | 1.84 | 8.33 | 2.34 | 1.22 | 2.98 | 3.58 | 3.55 | 1.44 | |
| | 1.05 | 2.14 | 1.77 | 1.67 | 1.31 | 0.80 | 1.43 | 1.52 | 2.10 | 0.30 | 2.10 | 3.51 | 3.53 | 0.48 | 3.81 | 2.16 | 2.66 | 3.65 | 1.67 | 1.66 | 5.52 | 2.75 | 1.06 | 3.12 | 2.84 | 2.04 | 1.52 | |
| | 0.94 | 1.85 | 1.48 | 2.30 | 1.32 | 0.36 | 1.23 | 1.99 | 1.80 | 0.55 | 1.95 | 3.67 | 4.17 | 0.38 | 4.01 | 2.00 | 2.32 | 4.66 | 1.75 | 1.46 | 5.51 | 1.94 | 0.90 | 4.00 | 2.29 | 2.11 | 1.49 | |
| | 1.71 | 2.44 | 1.82 | 2.98 | 2.00 | 1.46 | 1.90 | 1.28 | 2.89 | 0.23 | 2.00 | 4.96 | 4.07 | 0.06 | 4.58 | 1.70 | 3.06 | 6.60 | 1.58 | 1.65 | 6.28 | 5.44 | 1.51 | 3.27 | 4.23 | 2.26 | 1.03 | |
| Ventral Prefrontal Cortex | 1.49 | 2.64 | 1.90 | 2.25 | 2.25 | 0.65 | 1.81 | 2.89 | 2.65 | 0.51 | 2.65 | 4.54 | 5.61 | 1.05 | 4.37 | 2.10 | 3.39 | 5.40 | 1.41 | 1.82 | 6.07 | 4.62 | 2.09 | 4.49 | 3.13 | 1.98 | 0.98 | Ventral Prefrontal Cortex |
| | 1.87 | 3.05 | 1.62 | 1.38 | 2.87 | 0.60 | 1.29 | 1.68 | 2.17 | 0.59 | 2.34 | 5.14 | 5.19 | 0.60 | 4.90 | 2.97 | 3.51 | 5.77 | 1.18 | 2.56 | 6.22 | 3.76 | 1.55 | 4.34 | 4.45 | 1.98 | 1.84 | |
| | 1.77 | 3.23 | 1.95 | 3.24 | 1.63 | 1.70 | 1.93 | 2.05 | 3.09 | 0.97 | 2.63 | 4.89 | 5.68 | 0.46 | 4.76 | 3.37 | 3.14 | 4.83 | 2.45 | 2.65 | 6.96 | 2.43 | 1.30 | 3.31 | 4.42 | 2.14 | 1.39 | |
| | 1.55 | 2.48 | 1.82 | 2.09 | 1.58 | 0.96 | 1.64 | 2.04 | 2.27 | 0.69 | 2.61 | 3.87 | 3.08 | 0.59 | 3.31 | 2.29 | 2.90 | 4.06 | 1.91 | 1.96 | 5.43 | 2.95 | 1.24 | 3.40 | 2.54 | 2.18 | 1.61 | |
| Anterior Cingulate Cortex | 0.41 | 1.81 | 1.36 | 1.55 | 1.16 | 0.85 | 1.44 | 1.59 | 2.12 | 0.42 | 1.40 | 3.46 | 3.63 | 0.58 | 2.99 | 2.23 | 2.25 | 4.04 | 1.28 | 1.80 | 4.99 | 2.36 | 1.01 | 2.96 | 2.42 | 1.74 | 0.48 | Anterior Cingulate Cortex |
| | 0.63 | 1.51 | 1.12 | 1.17 | 1.21 | 0.65 | 1.27 | 1.45 | 1.72 | 0.30 | 1.70 | 2.97 | 3.07 | 0.59 | 2.32 | 1.68 | 2.01 | 3.54 | 1.07 | 1.53 | 4.35 | 1.92 | 1.23 | 2.27 | 1.87 | 1.57 | 1.05 | |
| | 1.49 | 1.92 | 1.39 | 1.22 | 1.02 | 0.70 | 1.31 | 1.89 | 1.67 | 0.57 | 1.93 | 3.23 | 3.38 | 0.50 | 2.41 | 1.67 | 2.21 | 3.52 | 1.62 | 1.51 | 4.46 | 2.13 | 0.82 | 2.61 | 2.13 | 1.70 | 1.85 | |
| | 1.62 | 2.18 | 1.38 | 1.49 | 1.36 | 0.76 | 1.29 | 1.49 | 1.83 | 0.58 | 2.05 | 3.15 | 3.18 | 0.53 | 2.61 | 2.22 | 2.36 | 3.08 | 1.69 | 1.65 | 5.03 | 2.45 | 1.14 | 2.62 | 2.42 | 1.56 | 1.50 | |
| Dorsal Premotor Cortex | 1.83 | 2.22 | 1.54 | 1.61 | 1.32 | 1.11 | 1.42 | 1.90 | 2.09 | 0.59 | 2.38 | 3.48 | 3.20 | 0.54 | 3.04 | 2.20 | 2.33 | 3.62 | 2.05 | 1.90 | 5.17 | 2.55 | 1.18 | 2.87 | 2.73 | 1.91 | 2.47 | Dorsal Premotor Cortex |
| | 2.00 | 2.02 | 1.44 | 1.74 | 1.30 | 0.83 | 1.54 | 1.81 | 2.16 | 0.67 | 2.40 | 3.86 | 3.44 | 0.64 | 3.01 | 2.53 | 2.79 | 4.48 | 2.04 | 2.62 | 4.64 | 2.94 | 1.13 | 3.15 | 3.07 | 2.34 | 2.13 | |
| | 2.05 | 2.46 | 1.87 | 2.06 | 1.60 | 1.10 | 1.65 | 2.17 | 2.08 | 0.75 | 2.40 | 3.56 | 2.82 | 0.68 | 3.28 | 2.51 | 2.82 | 3.75 | 1.70 | 1.71 | 5.30 | 2.89 | 1.18 | 3.48 | 2.70 | 2.17 | 3.87 | |
| | 1.67 | 2.20 | 1.34 | 2.30 | 1.51 | 1.18 | 1.41 | 1.36 | 1.94 | 0.91 | 3.10 | 3.55 | 3.01 | 0.43 | 3.06 | 1.86 | 2.30 | 3.47 | 1.49 | 1.88 | 5.43 | 3.27 | 1.28 | 3.48 | 2.94 | 1.59 | 2.48 | |
| Ventral Premotor Cortex | 2.28 | 2.59 | 1.81 | 1.97 | 1.47 | 0.94 | 1.42 | 1.86 | 1.95 | 0.51 | 2.43 | 3.33 | 2.75 | 0.54 | 2.88 | 1.99 | 2.76 | 3.45 | 1.52 | 1.66 | 4.92 | 2.87 | 1.11 | 2.96 | 2.43 | 1.85 | 2.29 | Ventral Premotor Cortex |
| | 1.21 | 2.78 | 1.97 | 2.11 | 1.71 | 1.10 | 1.77 | 2.01 | 2.32 | 0.52 | 2.91 | 4.07 | 3.00 | 0.90 | 3.57 | 2.28 | 2.50 | 3.68 | 1.64 | 2.16 | 6.18 | 3.78 | 1.44 | 3.58 | 2.67 | 2.86 | 2.03 | |
| Primary Motor Cortex | 2.07 | 2.44 | 1.64 | 1.69 | 1.29 | 0.90 | 1.42 | 1.66 | 2.04 | 0.58 | 2.50 | 3.25 | 2.53 | 0.55 | 2.98 | 1.90 | 2.59 | 3.13 | 1.69 | 1.69 | 5.24 | 2.64 | 1.12 | 3.35 | 2.45 | 1.83 | 3.43 | Primary Motor Cortex |
| | 1.13 | 2.33 | 1.50 | 2.01 | 1.37 | 0.82 | 1.56 | 2.01 | 2.22 | 0.69 | 2.36 | 3.51 | 3.06 | 0.53 | 2.79 | 1.78 | 1.86 | 3.06 | 1.73 | 1.66 | 5.64 | 2.65 | 1.18 | 3.13 | 2.74 | 1.60 | 2.45 | |
| | 1.05 | 2.18 | 1.44 | 2.22 | 1.68 | 1.37 | 1.97 | 1.72 | 2.32 | 1.74 | 2.82 | 2.98 | 3.32 | 0.62 | 2.91 | 1.79 | 1.57 | 2.78 | 1.71 | 1.60 | 5.00 | 2.77 | 1.33 | 3.97 | 3.08 | 2.18 | 1.96 | |
| | 1.16 | 2.62 | 1.38 | 1.81 | 2.28 | 1.17 | 1.61 | 1.41 | 1.96 | 1.14 | 2.46 | 3.86 | 3.95 | 1.08 | 3.32 | 2.00 | 3.35 | 2.93 | 1.67 | 1.53 | 5.40 | 2.98 | 1.20 | 3.65 | 3.04 | 2.23 | 2.59 | |
| | 1.47 | 2.25 | 1.32 | 2.10 | 1.80 | 1.28 | 1.76 | 1.36 | 2.34 | 0.71 | 2.55 | 3.32 | 3.94 | 0.80 | 2.94 | 1.77 | 1.66 | 3.28 | 1.78 | 1.60 | 5.40 | 2.94 | 1.12 | 3.40 | 3.23 | 2.01 | 2.41 | |
| | 1.39 | 2.30 | 1.38 | 1.79 | 1.15 | 0.83 | 1.62 | 1.95 | 1.80 | 0.64 | 2.72 | 3.43 | 2.34 | 0.62 | 2.75 | 1.54 | 1.43 | 2.48 | 1.53 | 1.53 | 5.22 | 2.87 | 0.95 | 3.21 | 2.81 | 1.61 | 6.83 | |
| Somatosensory Cortex | 1.43 | 2.67 | 1.43 | 2.16 | 1.72 | 1.00 | 1.66 | 1.87 | 2.13 | 0.54 | 2.84 | 3.84 | 2.82 | 0.77 | 3.09 | 2.37 | 2.15 | 3.58 | 1.84 | 2.08 | 5.63 | 3.13 | 1.18 | 3.45 | 2.87 | 2.19 | 4.00 | Somatosensory Cortex |
| | 1.33 | 3.38 | 1.68 | 2.26 | 1.81 | 1.18 | 1.88 | 1.84 | 1.97 | 0.50 | 2.91 | 4.72 | 3.39 | 0.82 | 3.13 | 2.36 | 2.91 | 3.98 | 1.47 | 2.27 | 7.05 | 2.79 | 1.44 | 3.88 | 3.56 | 2.49 | 2.21 | |
| Caudate | 0.91 | 2.70 | 1.70 | 2.11 | 3.10 | 2.87 | 2.16 | 4.12 | 2.48 | 1.81 | 1.78 | 4.48 | 3.08 | 3.04 | 3.50 | 2.63 | 4.67 | 5.85 | 1.92 | 2.09 | 5.40 | 4.64 | 2.25 | 3.79 | 3.19 | 2.11 | 3.84 | Caudate |
| | 1.63 | 2.56 | 0.98 | 2.32 | 2.75 | 1.04 | 2.34 | 1.84 | 1.87 | 0.32 | 3.07 | 4.83 | 3.26 | 2.03 | 2.69 | 2.65 | 3.62 | 4.59 | 2.64 | 1.54 | 4.12 | 3.13 | 1.35 | 3.88 | 2.14 | 2.38 | 1.99 | |
| | 1.27 | 2.43 | 1.81 | 2.32 | 3.38 | 3.30 | 2.60 | 4.75 | 2.37 | 1.91 | 3.51 | 5.20 | 3.97 | 1.68 | 4.61 | 3.94 | 3.71 | 4.80 | 2.51 | 2.67 | 7.86 | 3.87 | 2.36 | 5.52 | 3.19 | 2.49 | 1.78 | |
| | 1.17 | 2.50 | 2.24 | 1.99 | 3.71 | 2.97 | 2.67 | 3.95 | 2.74 | 2.03 | 3.65 | 6.93 | 3.14 | 1.57 | 3.32 | 3.22 | 3.78 | 5.08 | 2.07 | 3.15 | 7.91 | 4.80 | 2.19 | 6.13 | 2.82 | 3.03 | 0.86 | |
| Putamen | 1.77 | 2.27 | 2.02 | 3.85 | 4.00 | 3.89 | 2.42 | 4.63 | 2.43 | 3.02 | 3.51 | 4.57 | 3.77 | 2.67 | 3.63 | 4.13 | 3.40 | 5.50 | 3.17 | 3.21 | 4.80 | 4.34 | 2.64 | 4.75 | 3.95 | 4.85 | 0.45 | Putamen |
| | 2.04 | 2.64 | 2.35 | 3.80 | 4.19 | 5.00 | 2.37 | 5.74 | 2.60 | 3.40 | 3.88 | 4.35 | 3.03 | 2.73 | 3.00 | 3.96 | 3.40 | 4.73 | 3.29 | 3.10 | 3.71 | 3.97 | 2.66 | 3.95 | 3.25 | 3.84 | 0.57 | |
| | 0.93 | 2.60 | 1.64 | 3.95 | 2.74 | 1.34 | 2.35 | 2.63 | 2.02 | 0.93 | 3.89 | 6.35 | 4.47 | 1.02 | 5.71 | 3.73 | 5.90 | 5.40 | 3.06 | 3.07 | 8.60 | 4.64 | 2.22 | 5.35 | 3.71 | 4.43 | 0.23 | |
| | 1.62 | 3.86 | 2.15 | 3.41 | 2.33 | 0.82 | 2.08 | 2.66 | 1.75 | 0.59 | 4.10 | 6.41 | 4.57 | 0.63 | 6.93 | 3.31 | 7.24 | 4.64 | 3.94 | 2.60 | 9.70 | 4.13 | 1.98 | 4.36 | 3.39 | 3.66 | 0.15 | |
| | 1.04 | 3.00 | 1.82 | 3.51 | 2.20 | 1.30 | 2.65 | 2.47 | 2.24 | 0.85 | 4.79 | 7.24 | 3.28 | 1.07 | 4.47 | 2.97 | 4.74 | 4.39 | 2.57 | 3.56 | 9.51 | 5.71 | 2.60 | 6.49 | 3.27 | 4.46 | 0.24 | |
| | 1.17 | 2.62 | 1.83 | 2.60 | 1.64 | 1.74 | 2.77 | 2.56 | 2.25 | 1.15 | 4.68 | 6.79 | 3.87 | 1.33 | 4.83 | 3.24 | 2.90 | 4.75 | 2.76 | 3.65 | 9.35 | 6.10 | 2.62 | 6.66 | 4.26 | 4.68 | 0.30 | |
| | 1.37 | 3.13 | 1.86 | 2.96 | 1.47 | 1.29 | 2.39 | 2.11 | 1.71 | 0.82 | 4.50 | 5.87 | 5.93 | 0.88 | 6.52 | 3.18 | 3.33 | 6.33 | 3.35 | 2.94 | 9.21 | 5.12 | 2.02 | 4.84 | 4.99 | 3.73 | 0.25 | |
| External Globus Pallidus | 1.42 | 2.70 | 1.94 | 2.11 | 2.45 | 2.67 | 2.92 | 3.24 | 2.88 | 1.57 | 4.10 | 6.69 | 3.36 | 2.50 | 4.13 | 3.19 | 1.95 | 5.11 | 2.09 | 4.01 | 7.88 | 5.34 | 2.42 | 6.38 | 3.70 | 4.94 | 0.38 | External Globus Pallidus |
| | 1.46 | 3.10 | 2.22 | 2.32 | 2.60 | 2.59 | 2.75 | 3.21 | 2.82 | 1.39 | 4.16 | 6.85 | 2.97 | 2.61 | 3.63 | 2.93 | 2.09 | 4.55 | 1.98 | 3.96 | 8.30 | 5.26 | 2.29 | 6.13 | 3.23 | 6.10 | 0.44 | |
| Internal Globus Pallidus | 1.74 | 3.17 | 2.15 | 3.20 | 3.61 | 2.90 | 2.66 | 4.19 | 3.07 | 2.04 | 3.78 | 5.35 | 3.42 | 1.85 | 3.98 | 3.17 | 2.71 | 5.12 | 2.34 | 3.36 | 6.57 | 3.78 | 2.03 | 5.18 | 3.99 | 3.74 | 0.28 | Internal Globus Pallidus |
| | 1.38 | 2.63 | 1.69 | 2.94 | 2.89 | 2.77 | 2.76 | 3.11 | 2.96 | 1.36 | 3.26 | 5.09 | 3.23 | 1.95 | 3.23 | 3.11 | 2.05 | 6.24 | 1.93 | 3.29 | 5.19 | 3.32 | 1.65 | 4.93 | 3.70 | 4.13 | 0.35 | |
| Amygdala | 0.23 | 1.51 | 1.26 | 1.05 | 0.79 | 0.60 | 1.11 | 1.22 | 1.85 | 0.35 | 1.27 | 2.41 | 1.80 | 0.52 | 2.16 | 1.48 | 2.23 | 2.59 | 1.09 | 1.18 | 3.23 | 1.67 | 0.83 | 2.43 | 1.47 | 1.56 | 0.98 | Amygdala |
| Thalamus | 0.74 | 2.58 | 1.81 | 2.61 | 3.06 | 1.92 | 2.85 | 3.17 | 3.32 | 1.18 | 2.81 | 6.58 | 3.61 | 1.64 | 4.54 | 3.13 | 2.47 | 7.73 | 1.97 | 4.33 | 7.84 | 4.54 | 2.33 | 6.37 | 4.68 | 3.78 | 0.30 | Thalamus |
| | 0.86 | 2.79 | 1.98 | 3.09 | 2.22 | 1.70 | 2.58 | 2.17 | 3.25 | 1.02 | 2.91 | 6.62 | 3.70 | 1.29 | 5.01 | 3.79 | 2.44 | 7.54 | 2.32 | 3.37 | 8.46 | 4.18 | 2.22 | 6.05 | 4.65 | 4.67 | 0.24 | |
| | 1.00 | 3.21 | 2.31 | 2.61 | 2.30 | 1.63 | 2.70 | 2.54 | 3.50 | 0.94 | 3.43 | 6.89 | 3.41 | 1.35 | 5.49 | 3.51 | 2.33 | 7.10 | 2.07 | 3.96 | 8.55 | 4.41 | 2.27 | 6.95 | 4.56 | 4.12 | 0.25 | |
| Hippocampus | 1.56 | 2.53 | 2.01 | 2.49 | 1.37 | 1.70 | 2.79 | 2.19 | 2.02 | 0.45 | 4.08 | 7.87 | 2.93 | 1.33 | 4.31 | 2.14 | 2.81 | 3.98 | 1.53 | 4.10 | 9.98 | 5.28 | 1.70 | 6.62 | 3.35 | 5.03 | 0.54 | Hippocampus |
| | 1.06 | 3.38 | 1.97 | 2.50 | 2.14 | 1.50 | 3.15 | 1.42 | 2.26 | 0.92 | 2.88 | 6.47 | 3.50 | 1.83 | 4.74 | 2.56 | 2.47 | 4.71 | 1.80 | 3.48 | 8.94 | 4.87 | 2.42 | 5.07 | 3.64 | 4.25 | 0.40 | |
| Substantia Nigra | 1.46 | 5.97 | 2.64 | 3.67 | 1.94 | 1.16 | 2.61 | 1.45 | 2.56 | 0.38 | 3.29 | 5.64 | 2.73 | 0.67 | 4.58 | 2.55 | 1.86 | 5.85 | 1.53 | 2.84 | 6.78 | 3.91 | 1.82 | 4.19 | 3.76 | 6.15 | 0.70 | Substantia Nigra |
| | 1.43 | 6.81 | 3.49 | 3.99 | 1.58 | 1.04 | 2.94 | 2.17 | 2.89 | 0.39 | 4.41 | 5.77 | 3.41 | 0.69 | 5.64 | 2.49 | 1.10 | 5.98 | 2.03 | 3.13 | 7.31 | 5.08 | 2.40 | 3.89 | 4.41 | 6.42 | 0.24 | |

0 — 7 — 14

*Barcode variant abundance (%)*

**Figure S4. Cell-type-specific genes and cluster annotations for NHP putamen**

A

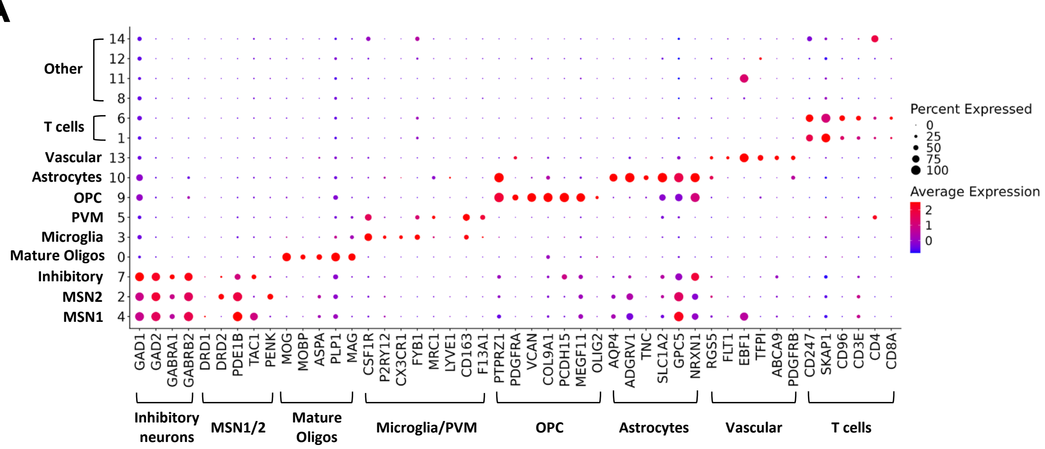


B

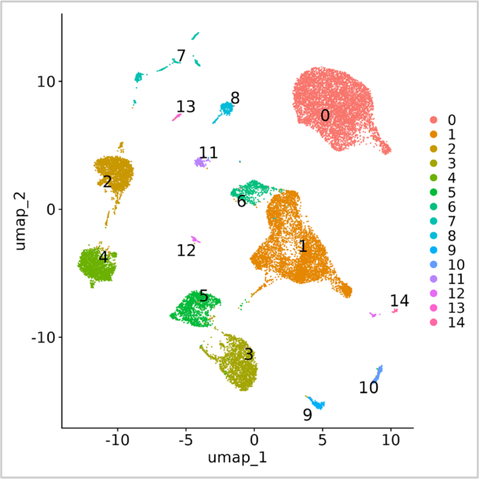

#### Figure S5. Cell-type-specific genes and cluster annotations for NHP substantia nigra

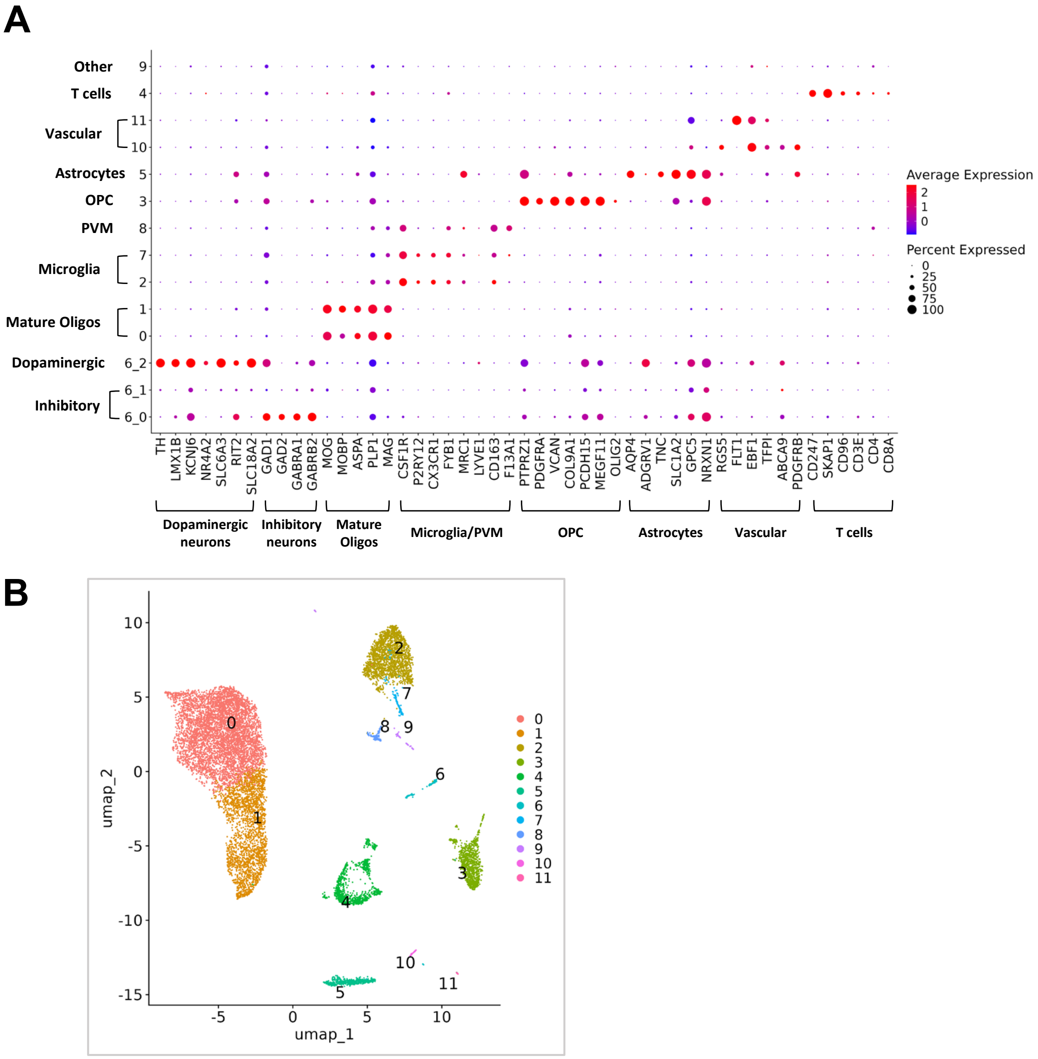

**Figure S6. Cell-type-specific genes and cluster annotations for NHP anterior cingulate cortex**

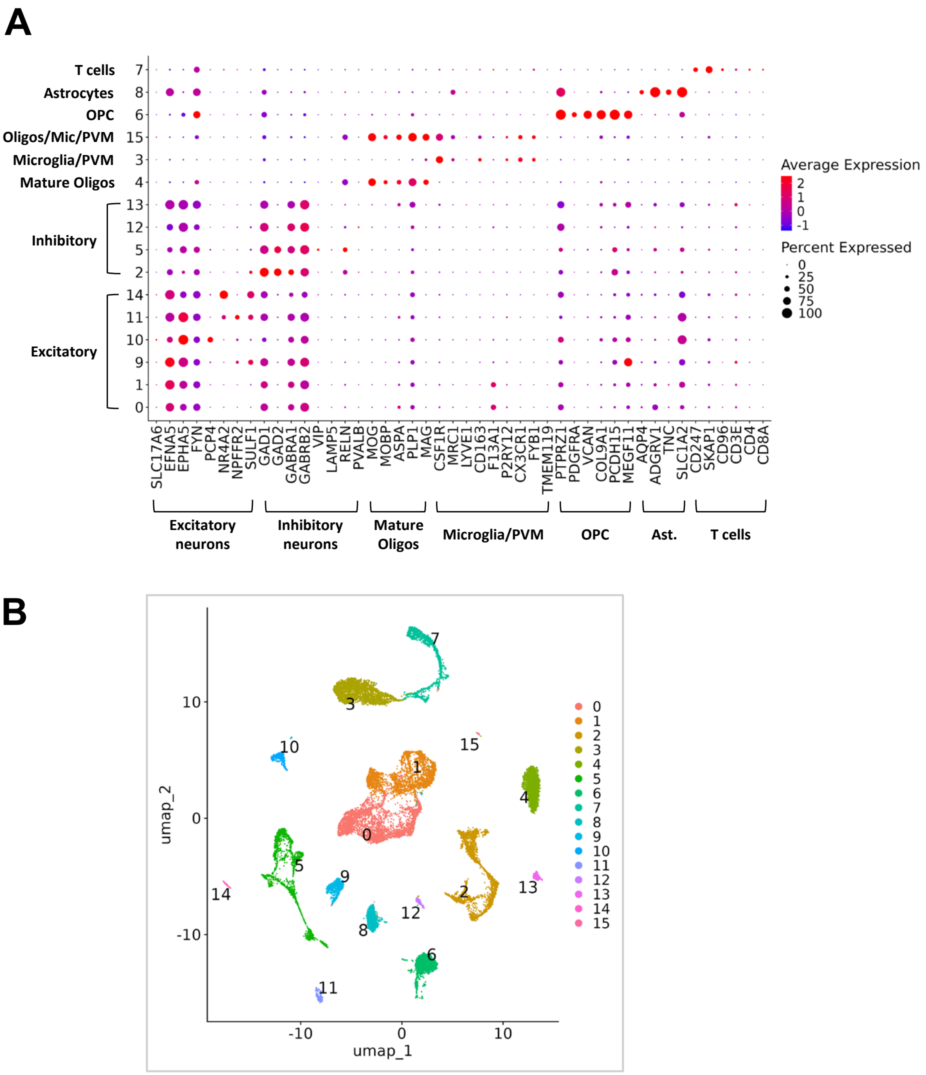

**Figure S7. Relative detection of transgene(+) nuclei via traditional 10x 3' GEX versus amplicon enrichment sequencing**

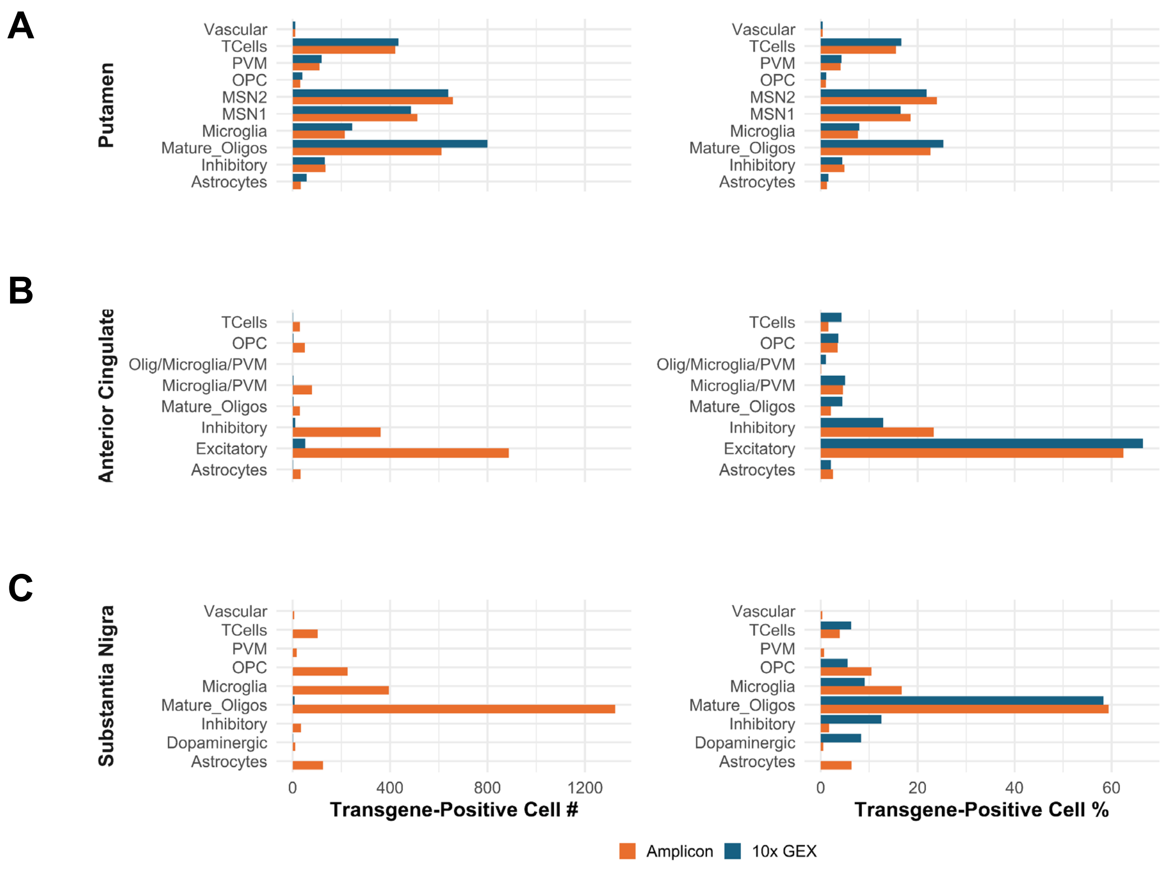
